# A trade-off between proliferation and defense in the fungal pathogen *Cryptococcus* at alkaline pH is controlled by the transcription factor GAT201

**DOI:** 10.1101/2023.06.14.543486

**Authors:** Elizabeth S. Hughes, Laura R. Tuck, Zhenzhen He, Elizabeth R. Ballou, Edward W.J. Wallace

## Abstract

*Cryptococcus* is a fungal pathogen whose virulence relies on proliferation in and dissemination to host sites, and on synthesis of a defensive yet metabolically costly polysaccharide capsule. Regulatory pathways required for *Cryptococcus* virulence include a GATA-like transcription factor, Gat201, that regulates Cryptococcal virulence in both capsule-dependent and capsule-independent ways. Here we show that Gat201 is part of a negative regulatory pathway that limits fungal survival at alkaline pH. RNA-seq analysis found strong induction of *GAT201* expression within minutes of transfer to RPMI media at alkaline pH. Microscopy, growth curves, and colony forming unit assays show that in RPMI at alkaline pH wild-type *Cryptococcus neoformans* yeast cells produce capsule but do not bud or maintain viability, while *gat201Δ* cells make buds and maintain viability, yet fail to produce capsule. *GAT201* is required for transcriptional upregulation of a specific set of genes, the majority of which are direct Gat201 targets. Evolutionary analysis shows that Gat201 is in a subfamily of GATA-like transcription factors that is conserved within pathogenic fungi but absent in model yeasts. This work identifies the Gat201 pathway as controlling a trade-off between proliferation and production of defensive capsule. The assays established here will allow characterisation of the mechanisms of action of the Gat201 pathway. Together, our findings urge improved understanding of the regulation of proliferation as a driver of fungal pathogenesis.

**Author Summary:** Micro-organisms face trade-offs in adapting to their environments. For example, pathogens adapting to host niches must balance investing in proliferation – reproduction and growth – against investing in defense against the host immune system. *Cryptococcus neoformans* is an encapsulated fungal pathogen that can infect human airways and, in immunocompromised people, can move to the brain to cause life-threatening meningitis. It is well appreciated that fungal persistence in these sites depends on production of a sugar capsule that surrounds the cell, hiding it from host detection. However, in both the lung and brain, fungal proliferation through budding is also a major driver of pathogenesis: both cryptococcal pneumonia and meningitis are characterised by high yeast burden. This presents a trade-off between production of a metabolically costly capsule and cellular proliferation. The regulators of *Cryptococcus* proliferation are poorly understood, as they are distinct from other model yeasts at the level of cell cycle and morphogenesis. In this work, we study this trade off growing *Cryptococcus* under conditions that approximate the alkaline surface of human airways, and that restrict fungal growth. We identify a GATA-like transcription factor, Gat201, and its target, Gat204, that positively regulate capsule production and negatively regulate proliferation. The GAT201 pathway is conserved within pathogenic fungi but lost in other model yeasts. Together our findings reveal how a fungal pathogen regulates the balance between defense and proliferation and highlight the need for improved understanding of proliferation in non-model systems.

## Introduction

*Cryptococcus neoformans* is an environmental saprophyte and a critical priority global human pathogen (World Health Organization 2022) that is a leading cause of death in HIV-positive individuals (Rajasingham et al. 2022; Park et al. 2009). *C. neoformans* is a basidiomycete fungus that is associated worldwide with bird guano and arboreal habitats (Nielsen, De Obaldia, and Heitman 2007; Springer, Mohan, and Heitman 2017; Litvintseva et al. 2011). *C. neoformans* can become an opportunistic pathogen following inhalation of airborne spores or desiccated yeast cells into mammalian airways (Walsh et al. 2019; Velagapudi et al. 2009; Powell et al. 1972; Giles et al. 2009). These infectious particles are initially metabolically inactive and must reactivate proliferation and simultaneously escape the immune system to infect host airways (Velagapudi et al. 2009; Ballou and Johnston 2017; Botts and Hull 2010; May et al. 2016). Proliferation of *Cryptococcus* in the upper airways or lung can be followed by dissemination through other host niches including to the central nervous system, causing fatal meningitis. Thus, both adaptation to diverse environmental niches and immune evasion are crucial for Cryptococcal virulence (Bose et al. 2003; Williamson 1997; A. Casadevall, Rosas, and Nosanchuk 2000; John R. Perfect 2006; Arturo Casadevall, Steenbergen, and Nosanchuk 2003).

However, proliferation and immune evasion can place competing metabolic demands on the fungal cell (J. Kronstad et al. 2012). For example, in the host, *Cryptococcus* cells produce a protective polysaccharide capsule. This unique and metabolically costly host evasion mechanism is dispensable for fungal proliferation and morphogenesis (García-Rodas et al. 2014), but is essential to *Cryptococcus* survival and dissemination in the host (O’Meara and Alspaugh 2012; Bose et al. 2003). By investigating conditions in which *C. neoformans* can either proliferate or protect itself, we can better understand the regulatory pathways controlling these competing demands.

We set out to probe this competition between proliferation and evasion during the initial switch to host-like conditions. Here, we reasoned that an informative model of initial infection would involve starting with stationary phase yeast cells that were then cultivated in “host-like” growth conditions, where different host-relevant conditions might give different insights into *Cryptococcus* adaptation. As the human airway surface becomes alkaline during each inhaled breath (Dusik Kim et al. 2021), we therefore investigated which genes are induced in *C. neoformans* stationary phase yeast cells directly after inoculation into *in vitro* conditions including host-like RPMI-1640 media at alkaline pH (RPMI). Our RNA-seq revealed strong induction of the *GAT201* pathway in these conditions.

The GATA-like zinc finger transcription factor Gat201 is a key regulator of virulence that acts in capsule-dependent (Liu et al. 2008) and capsule-independent ways (Chun, Brown, and Madhani 2011). *C. neoformans* strains with *GAT201* deleted have lower virulence in mouse models of infection (Liu et al. 2008) and reduced capsule (Jung et al. 2015; Gish et al. 2016; Jang et al. 2022). *GAT201* deletion strains are also more readily taken up by mammalian macrophages independently of capsule production (Chun, Brown, and Madhani 2011). The genes targeted by Gat201p have been mapped by RNA-seq and ChIP-seq (Homer et al. 2016; Chun, Brown, and Madhani 2011), yet the nature of its capsule-independent contributions to virulence remain unexplained. One challenge is that *gat201Δ* exhibits relatively weak phenotypes in standard microbiological growth conditions, making it difficult to probe Gat201-regulated virulence (Jung et al. 2015).

We observed that WT cells inoculated into RPMI media at alkaline pH proliferated poorly, but that deletion of *GAT201* dramatically improved proliferation and long-term viability. This suggests that poor growth in RPMI conditions is a consequence of regulated gene expression, controlled by the transcription factor Gat201, rather than a physiological response to nutrient starvation. Through RNA-seq, we identify *GAT201*-dependent transcriptional signatures of this phenotype and demonstrate that it is activated to suppress proliferation under alkaline conditions, independently of serum and of cyclic AMP. Correlating transcription factor activation to different microbial phenotypes can reveal signaling pathways required for adaptation to specific environments. Our findings of novel microbiological growth phenotypes dependent on *GAT201* will enable future mechanistic studies of this virulence pathway.

## Results

### *Cryptococcus neoformans* rapidly induces media-specific growth programs upon reactivation from stationary phase

We set out to determine which gene expression pathways are induced in *C. neoformans* stationary yeast cells when they reactivate in nutrient rich growth medium (YPD) versus host-like medium (RPMI-1640 + 10% Heat inactivated fetal calf serum; abbreviated RPMI) at 2 different temperatures (25°C or 37°C), and grown at 60 rpm shaking to mimic low-oxygen conditions. To investigate the early events of cellular reactivation, we used stationary phase cells incubated for 5 days in YPD at 30°C (Fig 1A). We sampled 2 biological replicates of cells prior to inoculation (0 minutes), and then sampled the cultures at 10, 30, 60, and 120 minutes following inoculation into either YPD or RPMI media at the two temperatures. We also collected a standard growth condition, mid-exponential phase cells reinoculated 1:30 into fresh YPD at 30°C for 180 min. Cells were examined for morphology (Fig 1B) and for transcriptional changes via RNA-seq (Fig 1C,D,E)

**Fig 1:**
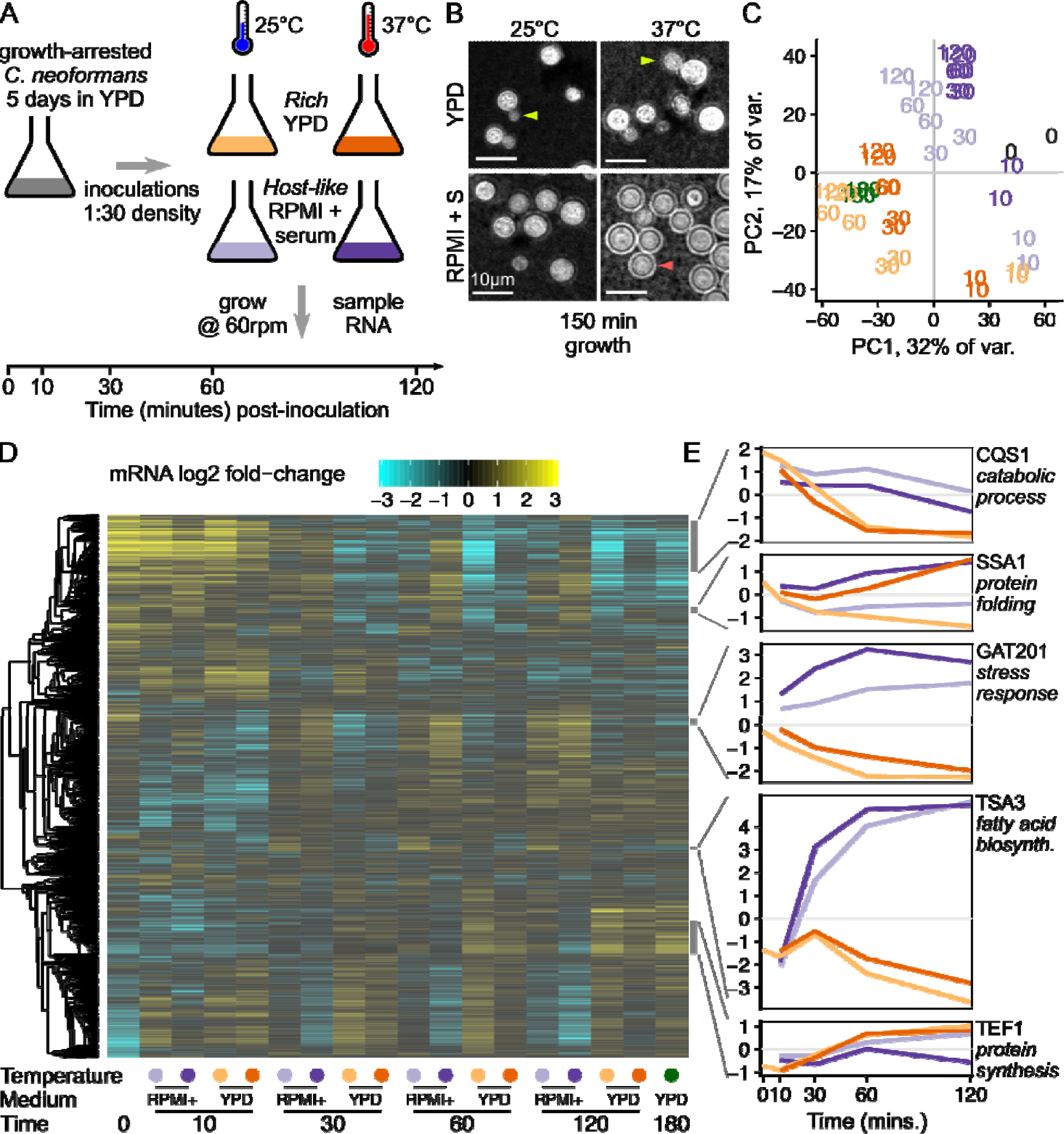
*Cryptococcus neoformans* rapidly induces media-specific growth programs upon reactivation from stationary phase. A, Design of time-course experiment to measure the contribution of media and temperature on reactivation of stationary phase cells. B, Budding (yellow-green arrow) is seen in YPD at both temperatures (top row) while capsule (light red arrow) is primarily seen in RPMI + serum at 37°C (bottom right). Micrograph shows India Ink staining of cells taken from one replicate of cells used for the RNA-seq experiment, 150 minutes after inoculation. C, Time from inoculation and media dominate the overall variance in gene expression, shown by principal component analysis on the regularized log-counts per gene in every replicate. Each replicate is plotted and labeled by the timepoint, with colours as in Fig. 1A (grey - 0 min, yellow - YPD 25°C, orange - YPD 37°C, light purple - RPMI 25°C, dark purple - RPMI 37°C), additionally with dark green for exponential phase rich media 30°C. D, Distinct clusters of co-regulated genes respond to reactivation in different media, shown by clustered heatmap of log2 fold-change per gene calculated by DESeq2. Each row represents an individual gene and rows are clustered by co-expression patterns (see methods), while each column represents the regularised log2 fold-change estimate across both replicates in a single condition. E, Representative genes from different clusters show distinct expression patterns, again in regularised log2 fold-change per gene. Colours of growth conditions are as in Fig. 1A.

We observed that cells reactivating in rich medium resumed growth and cell division, with visible buds 150 minutes after inoculation (Fig 1B). However, cells reactivating in RPMI medium showed no visible budding, and when grown at 37°C also produced a polysaccharide capsule visible with India Ink staining (Fig 1B). The RPMI media here is buffered with sodium bicarbonate, grown in a standard aerobic shaking incubator at 60 rpm. We observed the pH of the media rise to become alkaline over the timecourse, resembling human airway surface liquid that is alkaline during inhalation dependent on its bicarbonate content (Dusik Kim et al. 2021). We return to discuss the role of alkaline pH on the phenotypes later.

Consistent with observed changes in cell morphology, we identified distinct gene expression programs in the different media and temperatures. Principal component analysis (PCA) on the regularized log-counts per gene shows clear separation between growth conditions (Fig 1C and Fig S1). Principal components (PCs) 1 and 2 separate stationary phase cells and early timepoints (0 min, 10 mins) from later timepoints. PCs 1 and 2 also separate YPD media and RPMI media at later timepoints, while the exponential phase samples grown in rich media at 30°C group with the other rich media samples. PCA groups both biological replicates together for all sample conditions, indicating a highly reproducible experiment. PC3 separates samples grown at different temperatures with a clear divide between 25°C and 37°C (Fig S1). PC4 largely separates the 0 min samples from all later timepoints (Fig S1), emphasizing that induction of gene expression pathways is detectable after only 10 minutes in either growth medium and independent of temperature.

Clustering genes by their expression patterns reveals the combinatorial impact of media and temperature over time (Fig 1D). One cluster of 484 genes are highly expressed in stationary phase and decline in rich media, marked by *CQS1* that encodes *Cryptococcus* quorum sensing protein Qsp1 (Fig 1E, Table S1). *CQS1* is the most abundant transcript in cells in stationary phase conditions and RPMI media, as previously noted (Homer et al. 2016). This cluster of stationary-phase upregulated genes are also enriched in catabolic process functions (GO:0006091 generation of precursor metabolites and energy; GO:0005975 carbohydrate metabolic process), indicating that cells are starving.

A second cluster of 60 co-regulated genes are upregulated at 37°C in both media, marked by *SSA1,* encoding a heat shock protein (Fig 1E, Table S1). This cluster of genes is enriched in protein folding chaperones (GO:0006457, protein folding), including *HSP40, HSP60, HSP70, HSP90,* and *HSP104* family members, as expected from the conserved heat shock response that has been previously observed in *Cryptococcus* (Steen et al. 2002). A relatively small number of genes - 60 in this cluster - were unambiguously induced by increased temperature in both media (Table S1; and DGE analysis in Table S2).

A third cluster of 57 co-regulated genes is upregulated in RPMI compared to YPD, marked by the virulence-associated transcription factor *GAT201* (Fig 1E, Table S1). This cluster of genes is also enriched in associations with the stress response (GO:0006950, response to stress). These include cell wall-related enzymes (chitin synthase *CHS4*, chitinase *CHI2*, glucan glucosidase *EXG2*, Endoglucanase *LPI9*), as well as genes associated with redox metabolism (catalase *CAT2*, glutathione transferase CNAG_03848). Differential expression analysis of media conditions (Table S2) shows that *GAT201* is one of the most differentially expressed genes after 1 hour at both 37°C (∼24-fold, p < 10^−19^) and 25°C (∼13-fold, p < 10^−20^). Gat201 acts through other key transcription factors Gat204 and Liv3 (Homer et al. 2016). Consistent with *GAT201* induction 10 minutes after inoculation, we later observed induction of *GAT204* and *LIV3* expression in RPMI at 30 minutes and onwards (Fig S2).

A fourth cluster of 25 co-regulated genes is even more induced in RPMI, marked by *TSA3*, encoding a thiol-specific antioxidant protein, which is over 80-fold induced (Fig 1E). *TSA3* has been reported to be strikingly induced by temperature and hydrogen peroxide when grown in YNB media (Missall, Pusateri, and Lodge 2004). This cluster is enriched in genes involved in fatty acid biosynthesis (GO:0006629, lipid metabolic process).

Lastly, a large cluster of 321 genes associated with growth is induced in YPD at both temperatures and in RPMI at 25°C only, marked by translation elongation factor *TEF1* (Fig 1E). This cluster of genes is enriched for genes involved in protein synthesis, including ribosomal proteins (GO:0005840, ribosome), translation elongation factors, and amino acid production (GO:0006520, cellular amino acid metabolic process). The cluster is also enriched in genes involved in DNA segregation (GO:0000278, mitotic cell cycle) and mitochondrial biogenesis (GO:0005739, mitochondrion).

Overall, our data show that reactivating cells rapidly activate transcription and biosynthetic pathways, in distinct ways in different conditions. Cells induce both protein synthesis and cell cycle progression in YPD media, and proliferate by producing buds within 3 hours of inoculation. A different set of pathways are activated in RPMI-1640 + serum media, prominently including the *GAT201* transcription factor, genes associated with oxidative stress responses, and many genes of unknown function. In RPMI at 37°C, cells make a polysaccharide capsule that is associated with cellular defense, but do not start budding. This suggests that cells are making a condition-dependent decision between proliferation and defense, and that this may be achieved via differential expression of transcription factors.

### GAT201 determines cellular phenotypes during reactivation

We next tested the effect of *GAT201* on reactivation phenotypes, reasoning that the strong induction of *GAT201* and its transcriptional targets in RPMI made this pathway a good candidate regulator. We used two independently generated mutants: *gat201Δm* is a complete deletion of the reading frame from start codon to stop codon from the Madhani collection (Liu et al. 2008) and *gat201Δb* is a disruption of the protein from the Bahn collection (Jung et al. 2015).

We verified the mutations to the *GAT201* locus by PCR. We also made 2 independent complemented strains *GAT201-C1* and *GAT201-C2* by expressing *GAT201* from its native promoter and terminator at a genomic safe haven locus in the *gat201Δm* background (Figs 2A and 2E). We initially characterize these strains in RPMI-1640 medium without serum, and later return to examining the effects of serum.

**Fig 2:**
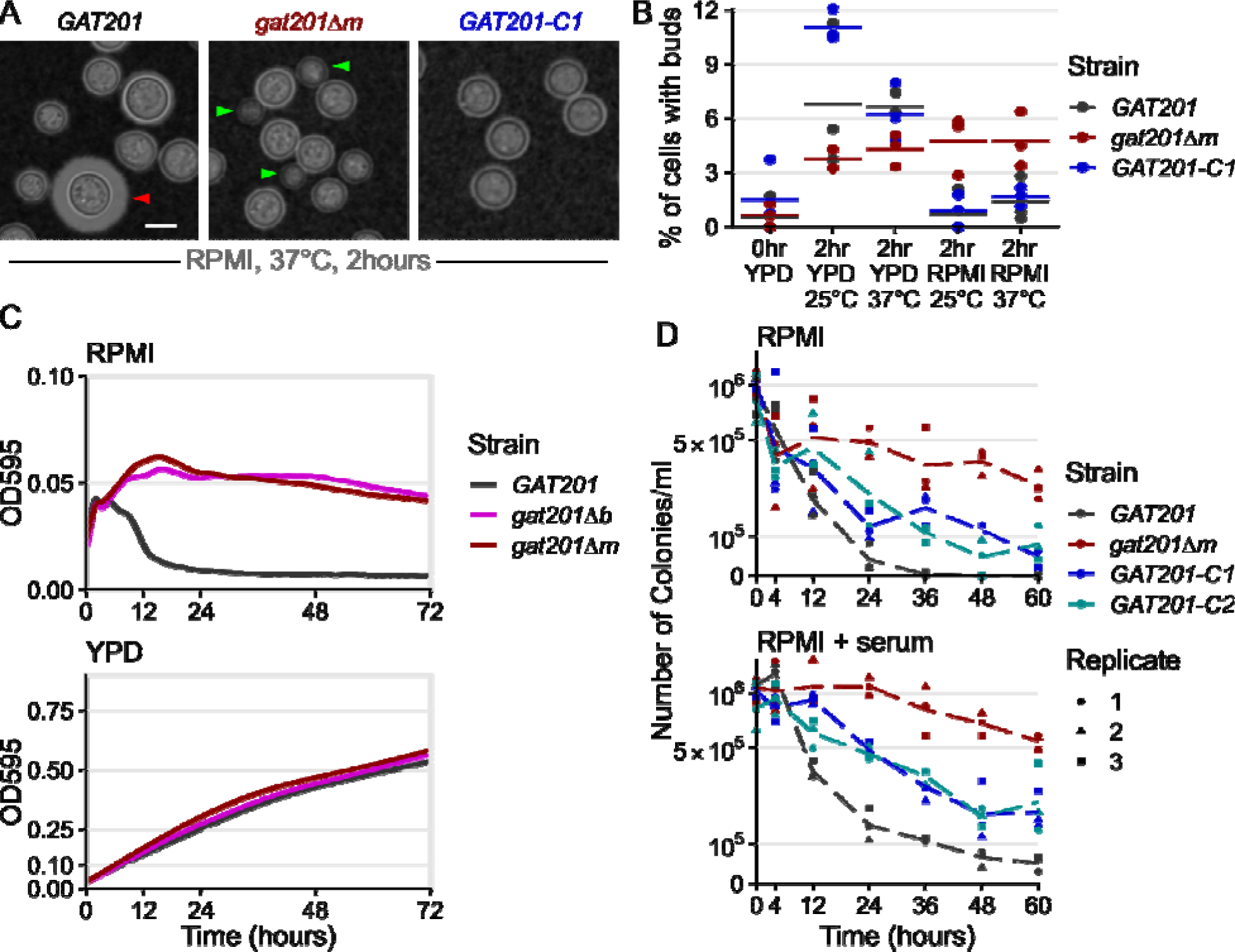
*GAT201* represses the proliferation and viability of *Cryptococcus neoformans* during reactivation in RPMI medium. A, *GAT201* promotes capsule biosynthesis and represses budding in RPMI-1640 medium (without serum) at 37°C 2 hours after inoculation. Micrographs show *GAT201* (H99), *gat201Δm*, and complemented *GAT201-C1* strains, stained with India Ink, capsule highlighted with red arrow and buds highlighted with green arrows. *GAT201-C1* complements the budding phenotype but does not clearly complement the capsule phenotype. B, Quantification of budding index at 2 hrs (% budded cells) shows that *gat201Δm* cells reactivate to produce buds in RPMI, (n= >100 cells per replicate, with 3 biological replicates per condition). Figs 2A and 2B are taken from the same experiment, and larger sets of representative cells are shown in Fig S3. C, *GAT201* (H99) cell populations reactivating in RPMI show a fall in density after 10 hours growth, which is absent in *gat201Δ* strains and absent during growth in rich YPD media. Growth curves of optical density at 595 nm (OD_595_) were collected via plate reader from 7 biological replicates, 3 technical replicates each, at 37°C. Note the different y-axis limits in the subpanels, reflecting higher final OD in rich media. D. *GAT201* (H99) cells reactivating in RPMI or RPMI + serum show a decline in viability after 12-24 hours, which is absent in *gat201Δ* and partially restored by complementing *GAT201*. The decline in viability is more severe in RPMI without serum than it is in RPMI with serum. Colony forming units per ml of culture were measured by serial dilution on plates, in 3 biological replicates; individual replicates are plotted as dots with a dotted line connecting the medians.

We observed that *GAT201* represses proliferation during reactivation in RPMI medium. 2 hours after inoculation in RPMI at 37°C, wild-type *GAT201* strains produce capsule and have few visible buds, while *gat201Δ* strains have visible buds and small capsule (Fig 2A, Fig S3). There is no difference in phenotype during growth in YPD (Fig S3). Genetic complementation of *GAT201* again represses bud formation, but only partially restores capsule. *GAT201* deletion mutants have previously been shown to be defective in capsule (Jang et al. 2022; Gish et al. 2016; Liu et al. 2008). The partial complementation of capsule in our strains with *GAT201* integrated at a genomic safe haven locus is likely to be due to the reduced expression of *GAT201* mRNA in this strain, measured by RT-qPCR as roughly ∼10x lower than in wild-type (Fig S4). This RT-qPCR also confirms the *GAT201*-dependent expression of *GAT204* and *LIV3*, which is partially restored in partially complemented strains and thus dependent on the abundance of *GAT201* (Fig S4).

Given the observed transcriptional changes within 10 minutes of reactivation, we quantified the budding index in the same samples at 2 hours post reactivation (Fig 2B), which is long enough for *C. neoformans* yeast to complete a single cell cycle (Yamaguchi et al. 2007). This early time point was chosen because it allowed detection of reactivation from growth without the possibility of measuring repeated budding or proliferating daughter cells. This confirmed that, upon reactivation from stationary phase, 6-10% of wild-type *GAT201* cells produce visible buds in rich media, but only 1-2% in RPMI media. Deletion of *GAT201* may reduce budding during reactivation in rich media, but increases budding in RPMI, with roughly 5% of *gat201Δ* cells budding within 2 hours, a more than 2-fold increase. Genetic complementation of *GAT201* reduces bud formation to near-wild-type levels.

Given this G*AT201*-dependent distinction in budding during reactivation, we investigated longer-term growth. While wild-type cells in RPMI medium did initially increase in optical density for 4 hours, density then rapidly declined within 10 hours (Fig 2C). In contrast, *gat201Δ* cultures for both mutants continually increased in cell density over 12 hours and maintained a higher OD_595_ of 0.05 over 3 days (Fig 2C). This effect is media-specific: *GAT201* cells and *gat201Δ* mutants grew similarly in rich media, reaching a similar OD_595_ of around 0.5 after 3 days (Fig 2C).

To determine if the *GAT201*-dependent reduced cell density in RPMI represented a loss of viability, we quantified colony-forming units (CFUs) from cells grown in RPMI at 37°C (Fig 2D). All cultures started with approximately the same number of stationary cells per ml (1×10^6^, OD_595_ = 0.1). By 24 hours the number of viable cells/ml in the *gat201Δ* mutant reduced by half, to approximately 5×10^5^, compared to a 10-fold reduction to 4×10^4^ in the wild-type, an order of magnitude less and a 25^th^ of the starting cell number. By 36 hours viability in the *gat201Δ* mutant strain had decreased by slightly more than a third of the starting cell number (3.7×10^5^ cells/ml) while *GAT201* wildtype viability was 2 orders of magnitude less than the starting cell number (1×10^4^ cells/ml). *GAT201* viability dropped to zero by 48 hours, indicating these cells are inviable following growth in RPMI media at 37°C after 2 days. In contrast, the *gat201Δ* mutant strain remained viable for up to 60 hours post-inoculation, with just under a quarter of the starting cell number (2.7×10^5^ cells/ml) still viable. Genetic complementation of *GAT201* reduced viability, although not to wild-type levels.

We next asked if the addition of serum (a host-like component) would alter the *GAT201*-dependent loss of viability, we repeated the CFU assay in RPMI + 10% serum at 37°C (Fig 2D). A similar pattern to RPMI was observed in RPMI + serum, however the number of colonies/ml overall were greater in RPMI + serum in both strains. The order of magnitude difference was maintained between the wild-type and *gat201Δ* mutant strains, with 1/20^th^ and half (5×10^4^ and 5.4×10^5^ cells/ml) of the starting cell number, respectively, at 60 hours post inoculation into fresh media. This indicates that the addition of serum to the media confers an overall benefit to cell viability in these conditions (Fig 2D), but that the effect is independent of *GAT201* activity. Genetic complementation of *GAT201* again partially restored the mutant phenotype.

Together, our data demonstrate that during reactivation *GAT201* determines cellular phenotypes beyond capsule production. When grown in RPMI medium, *GAT201* cells produce few buds within the first 2 hours of reactivation, lose optical density by 10 hours, and lose viability after 12-24 hours. The striking order-of-magnitude differences in viability, and qualitative differences in optical density, at longer timepoints, are prefigured by over 2-fold differences in budding index at earlier timepoints. Again, these experiments were conducted in an RPMI formulation buffered with sodium bicarbonate, grown under aerobic conditions so that the pH rose to become alkaline over the course of the experiment. We return to test the role of pH later, but first address the role of serum and *GAT201* on gene expression.

### How does serum affect the phenotype and gene expression?

Given the differential viability of wild-type and *gat201Δ* cells in RPMI media both with and without serum, we investigated the transcriptional pathways that might be responsible for these phenotypes. We designed an RNA-seq experiment with 2 deletion strains (gat201Δm and gat201Δb) matched with two congenic wild-type *GAT201* strains (KN99 MATa and MATalpha) (Nielsen et al. 2003), each measured in 2 biological replicates. Thus, there are effectively 4 replicates per relevant genotype. We first checked the effects of serum on capsule and budding in *GAT201* (KN99alpha) and gat201Δm. Microscopy confirmed that the major phenotypes are serum-independent: *GAT201* cells produce capsule and few buds in both RPMI and RPMI + serum, while gat201Δm produce less capsule and more buds in both conditions (Fig 3A, Fig S5).

**Fig 3:**
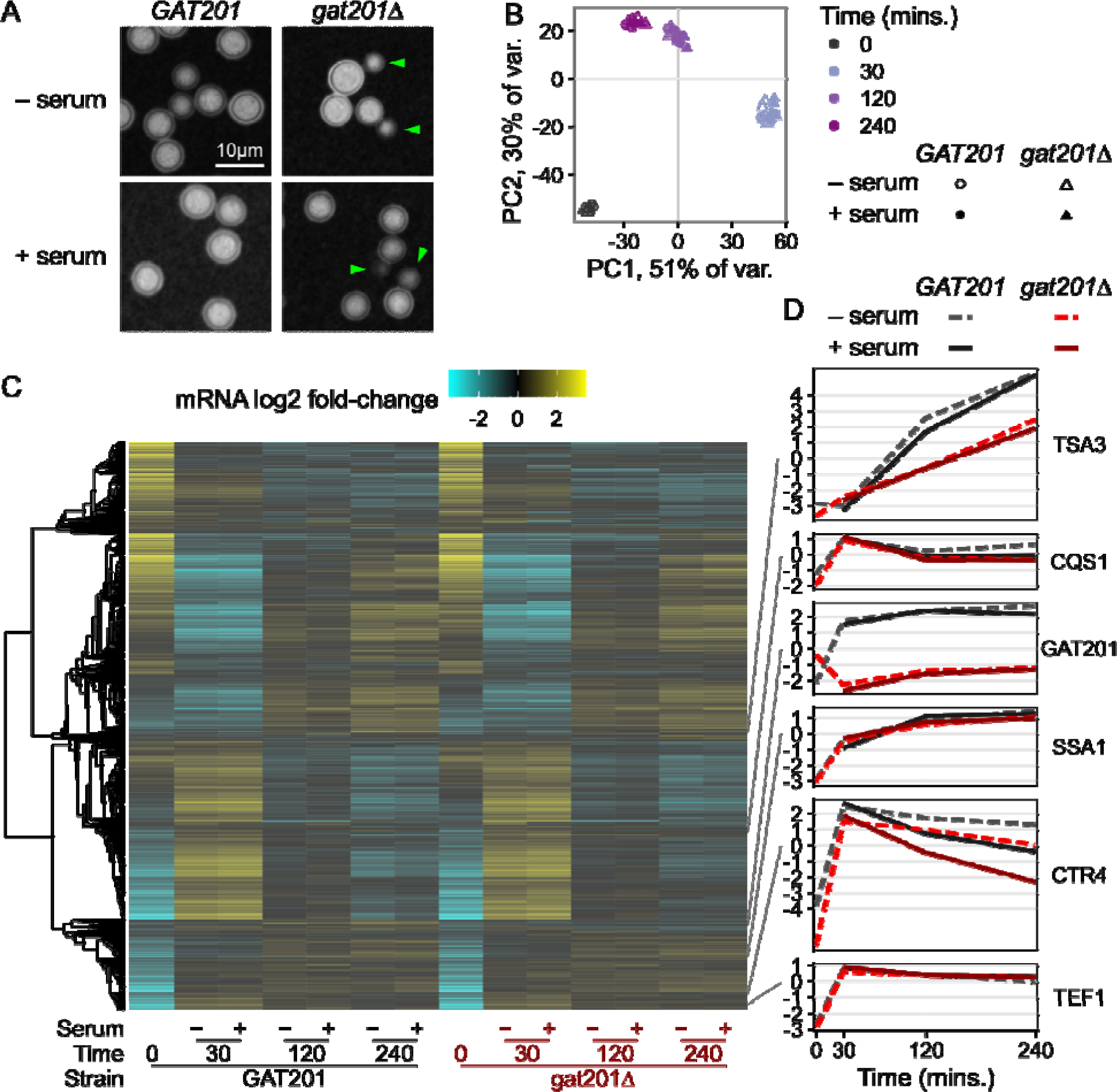
Serum is not the dominant driver of *GAT201*-dependent phenotypes in RPMI 1640 media. A, *GAT201* promotes capsule biosynthesis and represses budding in RPMI medium both with and without serum at 37°C, 2 hours after inoculation. Strains are *GAT201* (KN99alpha) and *gat201Δm*. B, Time from inoculation dominates the overall variance in gene expression regardless of serum addition or *GAT201* allele, shown by principal component analysis on the regularized log-counts per gene in every replicate. C, Only a small set of genes are differentially regulated by serum or by GAT201, shown by clustered heatmap of log2 fold-change per gene calculated by DESeq2. D, Representative genes show distinct expression patterns, again in log2 fold-change per gene.

We carried out RNA-seq of *GAT201* and *gat201Δ* strains reactivating, from stationary phase, in RPMI and RPMI + serum at 30 min, 2 hrs, and 4 hrs. Principal Component Analysis on the regularized log-counts per gene revealed that time after inoculation is the dominant driver of variance in this dataset (Fig 3B). 81% of variance is attributed to the first two principal components, in which datasets cluster together at each timepoint from all strains grown with and without serum. Principal component 3 (4% of variance) distinguishes between *GAT201* and *gat201Δ* strains (Fig S6), indicating that the RNA-seq detected a specific GAT201-dependent transcriptional program that we return to later. We were surprised not to detect a stronger serum-dependent effect, given previous work on the effect of serum on growth and capsule production in *Cryptococcus*, although this may reflect the short 4 hour time course (Zaragoza, Fries, and Casadevall 2003).

Consistent with serum not being the major driver of phenotype, we found few differences in the overall transcriptome-wide response to serum (Fig 3C, Table S3). Transcriptome-wide expression evolves over time with a large cluster of genes that are strongly upregulated at 30 minutes and still up at 2 and 4 hours. Conversely, a large cluster that is strongly downregulated at 30 minutes becomes less repressed at later timepoints. Expression of a set of representative genes selected from Fig 1 shows similar patterns to the previous dataset (Fig 3D). *TSA3* is strongly induced over time in RPMI and RPMI + serum, dependent on *GAT201* status. *CQS1* is highly expressed at 0 minutes and further induced in RPMI and RPMI + serum. *GAT201* itself is induced in RPMI and RPMI + serum in *GAT201* cells. The RNA-seq protocol used here is a 3’-end targeted assay that detects a transcript fragment of *GAT201* RNA in the two *gat201Δ* strains that have a truncated or deleted coding sequence; these transcript fragments do not encode a functional Gat201p. Both *SSA1* and *TEF1* are induced in RPMI and RPMI + serum as cells reactivate protein synthesis.

A small set of genes in wild-type cells differentially respond to the presence of serum at all time points. For example, copper transporter *CTR4* (Fig 3D) and copper starvation-induced membrane protein *BIM1* (Fig S7) are induced by 30 minutes growth in all media, and lower induction in media with added serum suggests differences in copper ion availability. Differential gene expression analysis with DESeq2 detected under 100 each of serum-upregulated and serum-downregulated genes at 5% FDR with at least 2-fold change, in *GAT201* strains (Fig S9, Table S3). Loss of *GAT201* dampened this response to serum, with about 4-fold more DEGs detected in *GAT201* than in *gat201Δ* (Fig S8). This dampened response in *gat201Δ* strains corroborates previous studies that found an association between expression of *GAT201* and downstream targets in alternative DMEM media with serum (Chun, Brown, and Madhani 2011).

Comparing our two RNA-seq datasets presented here indicates that the signature of *GAT201* pathway activation is consistent. Direct comparison of differential gene expression between the matched conditions of 0 minutes (pre-inoculation) and 2 hours RPMI + serum in wild-type cells shows moderate correlation, R = 0.27 (Fig S9, Table S2, Table S3). The major trends that are the focus of this study are consistent: *GAT201* and its targets are induced in RPMI + serum, as are many genes associated with protein synthesis. Despite this, there are some differences that may reflect local experimental conditions. For example, we observe that copper-dependent genes *CTR4* and *BIM1* are induced in RPMI in dataset 2 and not in dataset 1. This may reflect that the experiments were performed years apart by different experimentalists on different campuses. The datasets were also collected using different RNA-seq approaches. For dataset 1, we used RNATagSeq and rRNA depletion for full-length mRNA sequencing. For dataset 2, we used QuantSeq FWD for 3’-targeted mRNA sequencing.

### Most GAT201-dependent DEGs are direct targets of GAT201

A small and specific set of genes were found to be dependent on the expression of *GAT201*. After 5 days of growth in YPD (0 minute timepoint), there were only 82 2-fold differentially expressed genes at 5% FDR, up in wild-type *GAT201* strain compared to mutant gat201Δ. GO analysis suggested enrichment for upregulated genes involved in microtubule-based movement (GO:0007018), and lipid metabolism (GO: 0006629). 35 genes were down in *GAT201* compared to gat201Δ, with GO analysis highlighting probable changes in the cell wall (GO: 0016798, hydrolase activity acting on glycosyl bonds; GO:0005975, carbohydrate metabolic process) including the exoglucanase *EXG104*. After 30 minutes of incubation in RPMI or RPMI+serum, the *GAT201*-dependent differentially expressed genes are largely distinct from those observed in stationary phase. For example, *EXG104* is upregulated in *GAT201* after reactivation. The number of 2-fold differentially expressed genes at 5% FDR between *GAT201* and *gat201Δ* increases from 30 minutes up to 240 minutes (Fig 4A). We observed over 200 differential-expressed genes in each direction in RPMI (without serum) at 240 minutes (Fig 4B). Also, the magnitude of each gene’s differential expression between strains tends to increase over time (Fig 4C). Below, we focus on analysis of the RPMI 240 minutes timepoint.

Loss of activation of the Gat201 pathway in *gat201Δ* strains is confirmed by lower expression of known Gat201 targets during reactivation including *BLP1* (Fig 4C), *GAT204*, and *LIV3* (Fig S8). Functional enrichment analysis of genes up in *GAT201* cells highlighted changes at the cell surface, including transmembrane transport (GO:0055085) and cell-wall related terms carbohydrate metabolic processes (GO:0005975) and glycosyl hydrolase activity (GO:0016798). Metabolic pathway analysis highlighted *GAT201*-upregulated genes required for capsule biosynthesis (PWY-5114 PK MetaCyc, UDP-sugars interconversion), including UXS1/CNAG_03322 and UGD1/CNAG_04969 (O’Meara and Alspaugh 2012), consistent with the observed *GAT201*-dependent capsule synthesis during reactivation. Genes that are down in *GAT201* strains are enriched for ribosome biogenesis (GO:0042254) and related terms, indicating downregulation of the core biosynthetic process of protein synthesis. Genes down in *GAT201* strains were also enriched in carbohydrate metabolic processes (GO:0005975), indicating that different sets of carbohydrate-processing functions are upregulated and downregulated depending on *GAT201*. Overall, this implicates Gat201 in controlling carbohydrate metabolism and cell surface remodeling, and in repressing core biosynthetic processes related to growth.

We found that many of the *GAT201*-dependent genes are direct targets of Gat201. Previous measurements of binding of Gat201 to DNA by ChIP-seq found roughly 1200 enriched peaks out of approximately 6800 annotated Cryptococcal genes (Homer et al. 2016). We compared these targets to our list of differentially expressed genes at 240 minutes in RPMI (Fig 4D). Over 50% of the *GAT201*-upregulated genes here are direct targets (151/290), representing approximately 3x enrichment. The *GAT201*-downregulated genes at 240 minutes in RPMI are also about 2x enriched in direct targets (73/210). We see similar enrichment at earlier timepoints.

**Fig 4.**
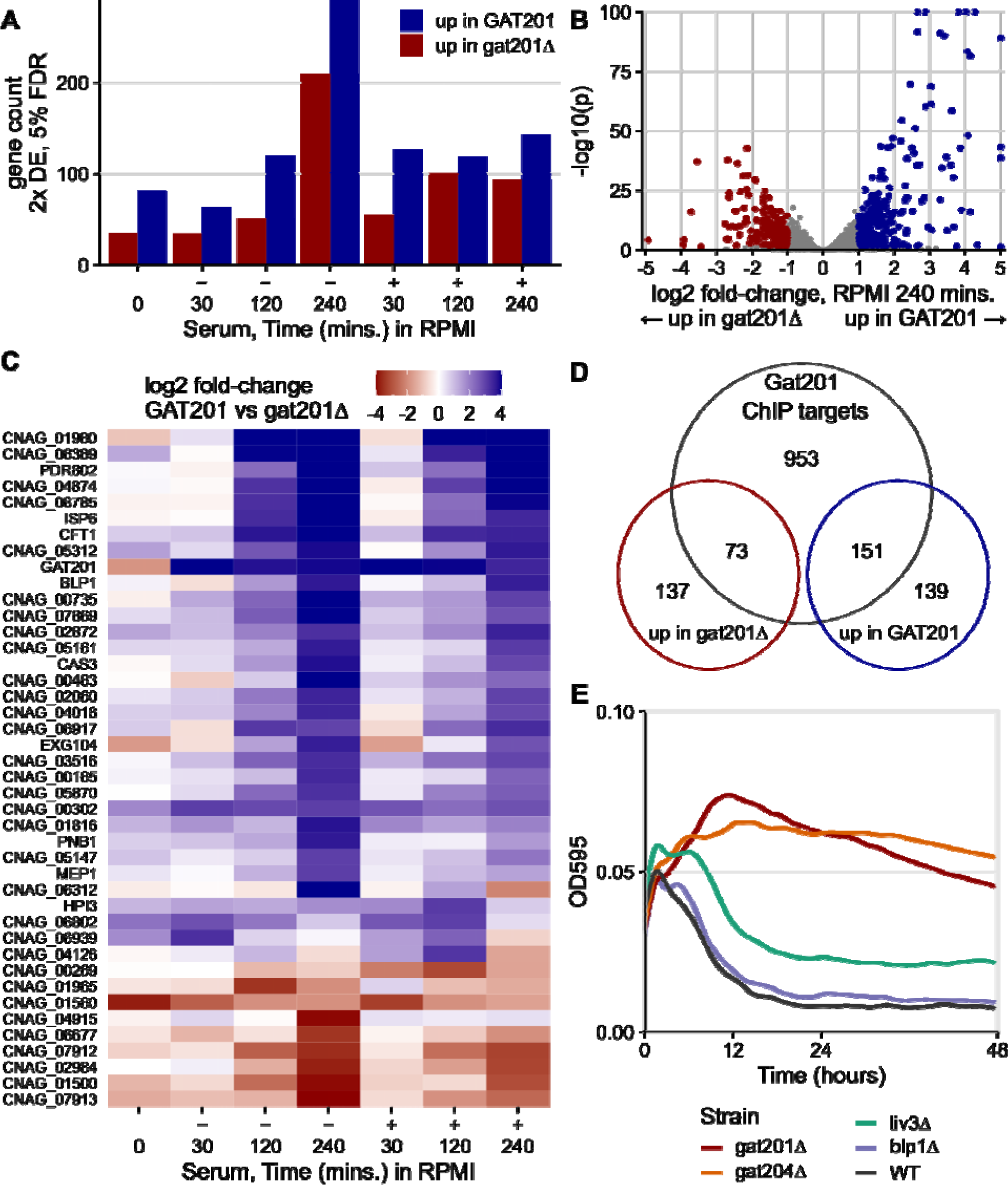
GAT201 specifically affects gene expression as cells reactivate, acting via its direct targets. A. There is more GAT201-dependent differential gene expression at later time points in activation. Fig 4A shows the number of 2-fold differentially expressed (DE) genes at 5% FDR at each combination of growth condition and time point. Differential expression is calculated by DESeq2 using the Wald test as the average over 4 samples: 2 wild-type and 2 deletion strains, each strain measured in biological duplicate. B. GAT201 promotes upregulation of specific genes more than downregulation. Volcano plot of log2 fold-change and p-value, with differential expressed genes calculated and coloured as in panel A. Genes with extreme p-values or fold-changes are plotted at the edge of the panel area. C. GAT201-dependent differential gene expression is more extreme at later timepoints. The panel shows all the genes that are at least 8x differentially expressed in any combination of condition and time, ordered by their average fold-change at 4 hours. D. Over half of the upregulated differentially expressed genes are direct targets of GAT201. Venn diagram shows the number DEGs in RPMI at 4 hours (as in panel B) compared to Gat201 targets measured by ChIP-seq from Homer et al. 2016. This is approximately a 3-fold enrichment. E. GAT204, as well as GAT201, is required to repress growth of cells in RPMI.

### *GAT204*, as well as *GAT201*, is required to repress growth of cells in RPMI

We next asked which *GAT201* targets could be required for the *GAT201*-dependent repression of proliferation and growth in RPMI. We took a reverse genetics approach focusing on phenotypically important *GAT201* targets and transcription factors, measuring growth curves of deletion mutants from the Madhani collection (Liu et al. 2008). Gat204 and Liv3 are transcription factors implicated in virulence that are Gat201 targets (Chun, Brown, and Madhani 2011) and whose direct target genes overlap with those of Gat201 (Homer et al. 2016). In RPMI medium, *gat204*Δ cells behave similarly to *gat201Δ* cells by increasing in density, unlike wild-type cells that decline in density within 10 hours (Fig 4E, Fig S10). Gat204 regulates approximately 30% of Gat201-regulated genes (Homer et al., 2016). An intermediate density is shown by *liv3*Δ cells (Fig 4E). Conversely, the barwin-like protein Blp1 that is required for the antiphagocytic function of Gat201 is dispensable for the growth phenotype: *blp1*Δ cells grow similarly to wild-type cells in RPMI (Fig 4E).

Of the genes that we have tested so far (Fig S10), only *GAT201, GAT204,* and *LIV3* are required for growth restriction in RPMI medium. Deletion of other *GAT201* targets, including transcription factors *PDR802* and *ECM2201*, and metalloproteinase *MEP1*, did not relieve restriction of growth (Fig S11).

Furthermore, we tested *GAT204* and *LIV3* genetic interactions in RPMI medium. The growth phenotypes of gat204Δ, liv3Δ and the double mutant gat204Δ liv3Δ were similar, all growing better than wild-type in RPMI but not as well as *gat201Δ* (Fig S11). This result is consistent with *GAT204* and *LIV3* operating in the same pathway, as previously reported. Overall, these data argue for the existence of a specific Gat201/Gat204/Liv3 dependent pathway that restricts growth in specific media at alkaline pH.

### GAT201 represses growth at alkaline pH but is required for growth in RPMI at neutral pH

We next examined the role of pH in determining the restricted growth phenotype. We had previously observed that, in RPMI buffered with sodium bicarbonate in our aerobic growth conditions, media alkalinization occurred regardless of the addition of serum or cells. However, in an alternative RPMI-like “CO_2_-independent media”, which is kept near neutral pH with a phosphate-based buffering agent, both *GAT201* and *gat201Δ* cells continue to increase in density over a 72 hour period (Fig S12). This differential growth in media with otherwise identical nutrient composition suggested that the buffering agent and/or pH was responsible for the phenotype.

Accordingly, we next isolated the effect of sodium bicarbonate (NaHCO_3_) on growth, by growing cells in RPMI with varying concentrations of NaHCO_3_ in aerobic conditions. With 24mM NaHCO_3_, the same as in the standard RPMI formulation used previously, we again observed that wild-type *GAT201* cells do not grow in the long-term but *gat201Δ* cells do grow (Fig 5, Fig S13). Surprisingly, at lower concentrations of NaHCO_3_ this effect changes, and in RPMI with no added NaHCO_3_ wild-type cells grow consistently while *gat201Δ* cells do not grow (Fig 5, Fig S13). Complementing *GAT201* into deletion strains restored the wild-type growth phenotype of growth at 0mM NaHCO3 and arrest at 24mM NaHCO_3_.

**Fig 5.**
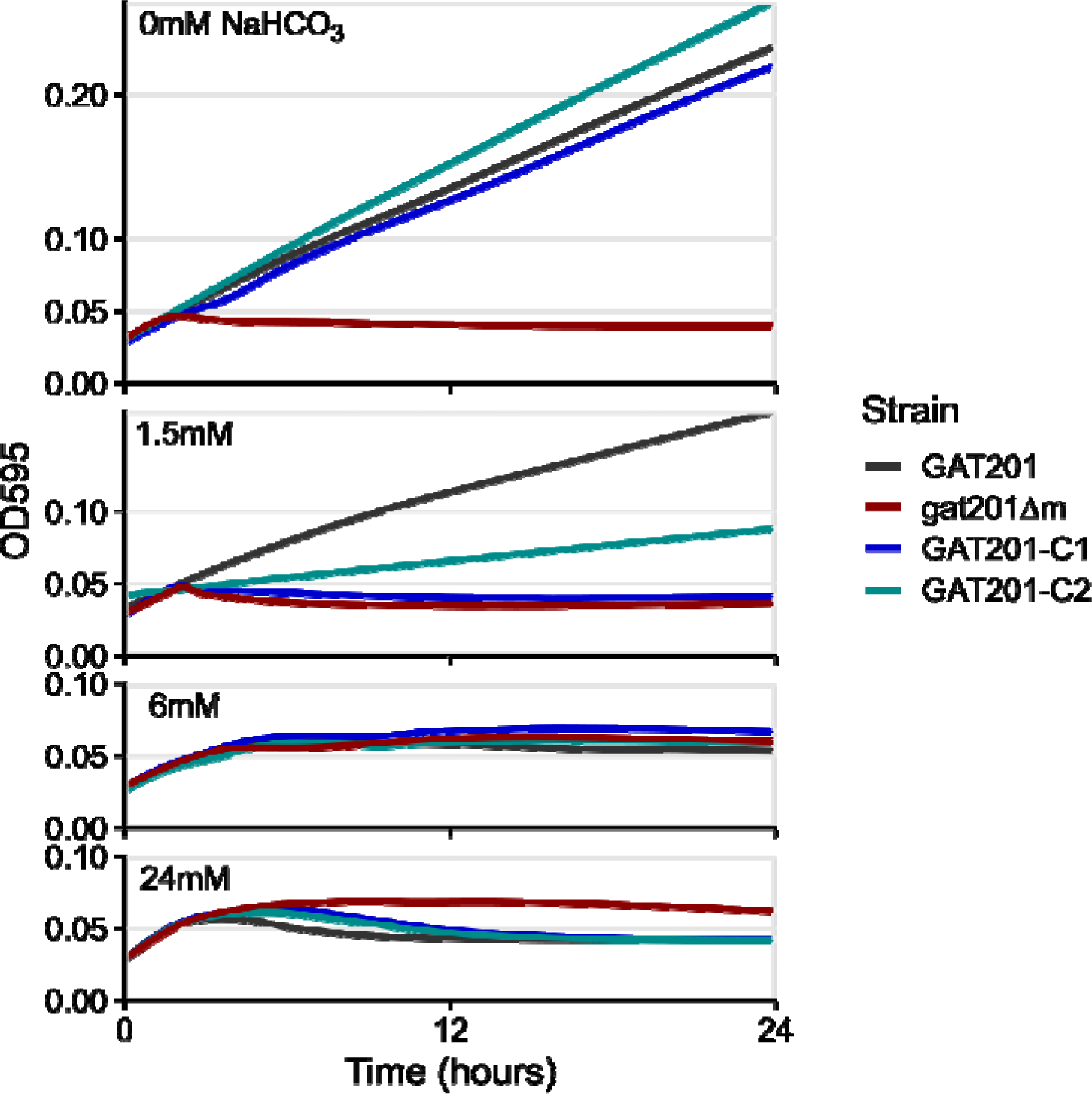
The effect of *GAT201* on growth depends on sodium bicarbonate (NaHCO_3_) Starting with an RPMI formulation lacking NaHCO_3_, we added either 0mM, 1.5mM, 6mM, or 24mM NaHCO_3_ and grew Cryptococcus for 24 hours. Wild-type *GAT201* cells grow in 0mM NaHCO_3_ but do not grow in 24 mM NaHCO_3_, while *gat201Δ* have opposite phenotypes of no growth in 0mM and growth at 24 mM. These cells have intermediate phenotypes at intermediate concentrations of NaHCO_3_, while complemented strains have growth phenotypes resembling wild-type. This figure shows the median of 3 technical replicates from a single biological replicate, and 2 further biological replicates are shown in Fig S13.

To interpret these data, note that bicarbonate ions in the media are in exchange with carbon dioxide in the air around the cultures (Michl, Park, and Swietach 2019). Addition of 24mM NaHCO_3_ leads to an equilibrium pH of about 7.5 in 5% CO2 (Michl, Park, and Swietach 2019), however at atmospheric CO_2_ of roughly 0.04% this media reaches pH ∼9.5 within hours. Without any added NaHCO_3_, the media pH remains close to neutral. Thus, the major differences in growth phenotypes that we observe occur after equilibration to more alkaline pH. Further work will be needed to dissect the effect of pH, buffers, and exogenous CO_2_ on *GAT201*-dependent growth, but the phenotype that *GAT201* promotes growth in RPMI media at near-neutral pH and represses growth at alkaline pH is reproducible in our hands.

Because cyclic-AMP dependent signaling through the Rim101 transcription factor also regulates growth at alkaline pH (Caza and Kronstad 2019; O’Meara et al. 2010), we tested if cyclic AMP signaling affects our observed *GAT201*-dependent phenotype. We found that addition of the exogenous cAMP does not substantially affect growth in RPMI with or without NaHCO_3_ added: again, wild-type *GAT201* cells grow far more than gat201Δm in the absence of NaHCO_3_, and the phenotype is reversed in 24 mM NaHCO_3_ (Fig S14). This shows that the *GAT201* pathway is largely independent of cAMP signaling, and thus of Rim101 signaling, indicating a distinct alkaline-responsive pathway controlled by *GAT201*.

### *C. neoformans* Gat201 is homologous to other GATA-family zinc finger proteins that regulate fungal growth and environmental responses

Lastly, we asked if Gat201 could be homologous to other fungal transcription factors that might indicate a conserved pathway. Gat201 is a 435 amino acid-long protein predicted to have only a single structured domain of 58 amino acids near the C-terminus, the GATA-like zinc finger domain (Fig 6A). This domain is found across a broad variety of transcription factors that integrate environmental signals and metabolism (Lowry and Atchley 2000). Searching for homologs of *C. neoformans* Gat201 by BLASTP (Sayers et al. 2022) detects many proteins with GATA-like domains and a variety of domain structures, consistent with the known diversity in GATA-like transcription factors (Lowry and Atchley 2000). Because there are multiple GATA-like domain proteins in each fungal species that we examined, we turned to more precise methods to search for true homologs of Gat201.

**Fig 6.**
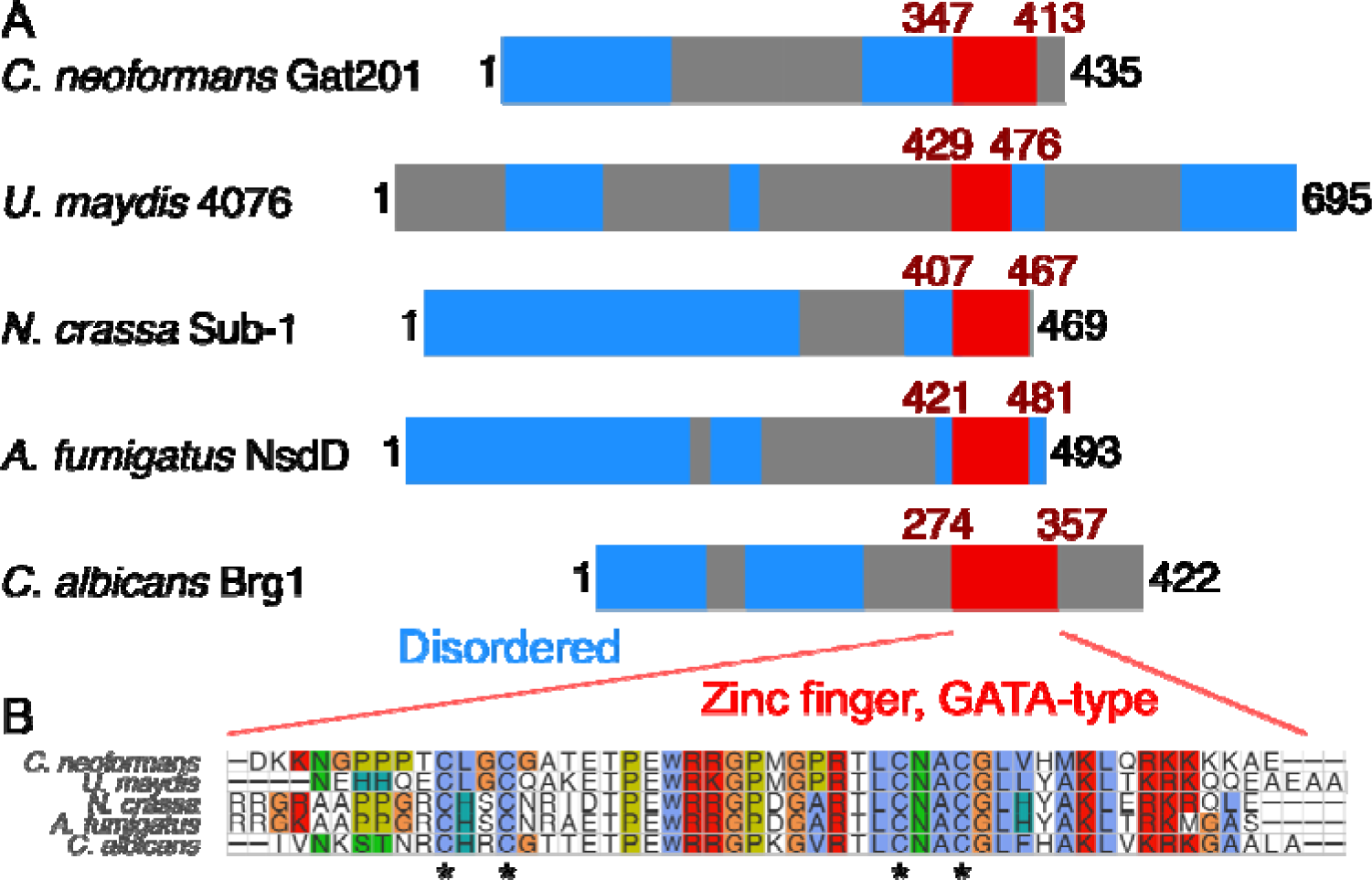
*C. neoformans* Gat201 is homologous to other GATA-family zinc finger proteins that regulate fungal growth and environmental responses A, Domain structure of Gat201 and 4 close homologs, with GATA-like zinc finger domain shown in red (Interpro IPR013088) and predicted unstructured regions in blue (MobiDB Lite consensus disorder), taken from Interpro (Blum et al. 2021). B, Multiple sequence alignment of the GATA-like zinc finger domains of homologs made with MUSCLE (Edgar 2004). Conserved cysteine residues typical of GATA-like zinc fingers are indicated with asterisks. An extended phylogeny and homology analysis is shown in Fig S15.

We turned to the PANTHERDB curated homology database, and selected the proteins closest to *C. deneoformans* Gat201 within family GATA transcription factor, PTHR45658 (Mi et al. 2019). This database groups Gat201 with fungal and amoebal proteins that have the same domain structure, including *S. cerevisiae* Gat2 and *C. neoformans* Gat204, as well as wc-2 transcription factors that are involved in light responses in filamentous fungi, that have an additional PAS sensing domain (Ballario et al. 1998). We then filtered to only homologs with the same domain structure as Gat201, performed a full-length protein alignment seeded with the GATA domain using MAFFT (Katoh and Standley 2013), and calculated a phylogenetic tree by Bayesian maximum likelihood using IQ-TREE (Minh et al. 2020).

Our analysis grouped *C. neoformans* Gat201 alongside a subset of other proteins with a single C-terminal GATA-like domain: *Ustilago maydis* UMAG_04076, *Neurospora crassa* Sub-1, *Aspergillus fumigatus* NsdD, and *Candida albicans* Brg1 (Fig 6A, Fig S15A). Several of these homologs have reported roles in regulating growth and environmental responses (Muñoz-Guzmán, Caballero, and Larrondo 2021; Lee et al. 2014; Cleary et al. 2012; Lu, Su, and Liu 2012). Fungal Gat201-like transcription factors are most closely related to amoeba homologs including gtaI, a GATA-family transcription factor that is required for morphological transitions in *Dictyostelium discoideum* (Katoh-Kurasawa et al. 2021). *S. cerevisiae* Gat2, Gat3, and Gat4 were grouped in a separate clade, while *C. neoformans* Gat204 was grouped into another clade close to a different set of amoebal homologs (Fig S15A). The zinc-finger domains of these Gat201 homologs are highly conserved (Fig 6B, Fig S15B), including the 4 cysteines that co-ordinate the zinc ion (Lowry and Atchley 2000). In addition, the AlphaFold2 protein structure database predicts a short unannotated alpha-helix-rich domain in *C. neoformans* Gat201, *N. crassa* Sub-1 and *A. fumigatus* NsdD (Varadi et al. 2022).

As an alternative method of searching for true homologs of *GAT201*, we looked for syntenic (gene-order conserved) homologs using the GenomicusFungi synteny browser (Nguyen et al. 2018). We searched against *C. deneoformans GAT201* (CNC06330), which is syntenic to *C. neoformans GAT201* across the locus. We found a set of basidiomycete syntenic homologs with a C-terminal GATA-like domain, including *U. maydis* UMAG_04076, that was identified as a close homolog in our phylogenetic analysis. Within the Agaricomycotina, the gene neighborhoods of these GATA-like domain proteins were dynamic, with extensive recombination. While there was limited preservation of wider gene synteny, GATA-like proteins within the Basidiomycota tend to be flanked upstream by genes encoding a Vms1-like protein (CNC06290, Ch3, UMAG_04964, Ch15), similar to *S. cerevisiae* YDR049W, (ChIV), and a lipid binding protein (CNC06300, Chr3; and UMAG_11290, Chr11) homologous to *S. cerevisiae* YJL036W (ChrX). There is limited longer range synteny downstream of *GAT201*, but the *GAT201* gene is closely flanked by a DEAD box RNA helicase (CNC06360, Chr3; and UMAG_04080, Chr11). *S. cerevisiae* encodes two orthologous genes of the DEAD-box RNA helicase: *DED1* and *DBP1* (YOR204W and YPL119C, located on Chr XV and Chr XVI respectively). Examination of synteny around these genes in *S. cerevisiae* identified genes that are well conserved in *Cryptococcus* and *Ustilago,* including *BEM3* (CNJ02560, Chr10; UMAG_11852, Chr2) and *HIS3* (CNH01620 Chr8; UMAG_11859 Chr2), neither of which were syntenic with GAT201 in either *Cryptococcus neoformans* or *Ustilago maydis,* nor even located on the same chromosome (GAT201: CNC06330, Chr 3; UMAG_04076, Chr 11). Overall, this analysis does not support a syntenic relationship between *GAT201* and other genes containing GATA-like domains beyond the Basidiomycota.

It remains possible that a more exhaustive analysis would detect more homologs or more precise recombination events involving related GATA-like domains. Short and less structured proteins like Gat201 contain less phylogenetic signal and are prone to “homology detection failure” (Weisman, Murray, and Eddy 2020). Still, the phylogenetic identification of a Gat201-like subfamily is supported by bootstrap analysis, by the similarity of the GATA domains, and by synteny analysis. Future work will be needed to assess which of Gat201’s predicted homologs have conserved molecular function or operate in a conserved pathway.

## Discussion

Proliferation is a major driver of *C. neoformans* pathogenesis: cryptococcosis pathology is driven by the accumulation of yeast in diverse host niches, and high fungal burden is a strong correlate of poor outcomes (J. W. Kronstad et al. 2011; Ballou and Johnston 2017; Brouwer et al. 2004; Bicanic et al. 2009). To proliferate in the host and cause disease, *C. neoformans* yeast must rapidly adapt to the lung environment, characterised by nutrient limitation, high temperature (37°C), and transient high pH (>8.5) (Dusik Kim et al. 2021). In this study we modeled the early events of the fungal transition from stationary phase to conditions that support proliferation (rich media; 25°C or 37°C, pH 5.5) and those that stimulate defense through the formation of a polysaccharide capsule (RPMI+serum; 37°C, pH 8.5-9). At early time points, we observed the activation of cryptic pathways in host-like media (RPMI-1640 with or without serum) that correlated with expression of the virulence-associated transcription factor *GAT201*. Surprisingly, activation of these pathways was associated with restriction in budding and, later, with loss of viability. By studying both short term and extended growth in alkaline conditions, we identified a cryptic growth program not revealed by previous work. Future work will leverage these conditions to reveal the molecular mechanisms underpinning Alkaline-Restricted Growth (ARG).

*C. neoformans* is known to be extremely sensitive to alkaline pH, failing to grow above pH 8.5 (Howard 1961; Levitz et al. 1997). However, it is surprising that this alkaline-restricted growth phenotype is rescued by deletion of the *GAT201* transcription factor, and that the same transcription factor blocks growth at neutral pH in otherwise identical media. The fact that deletion of a transcription factor restores budding, density, and viability indicates that the observed restricted growth of wild-type cells is a consequence of a regulated gene expression program. This genetic constraint on growth is not solely a physiological constraint due to alkaline pH or lack of nutrients.

Previous studies in *C. neoformans* have focused on genes whose loss further restricts growth at alkaline pH (Ost et al. 2015). Here, by contrast, we demonstrated that loss of the Gat201 pathway enables growth at alkaline pH. This restriction is apparent as early as 2 hours, when there is little yeast bud emergence and production of substantial polysaccharide capsule in wild-type cells. Deletion of *GAT201* restores yeast cell budding and also disrupts the production of capsule, and this is consistent with microscopy data published in a previous study that focused on capsule rather than on proliferation (Gish et al. 2016). From 12 hours onwards, we found that wild-type *C. neoformans* becomes inviable, while viability was rescued by deletion of *GAT201*. These assays for Gat201 pathway activation will enable dissection of the pathway and its involvement in proliferation, that could shed light on Gat201’s role in promoting virulence.

We observed transcriptional activation of *GAT201* within minutes of reactivation of *C. neoformans* in RPMI, independent of serum and temperature. Correspondingly, in these conditions we observed upregulation of many direct targets of *GAT201*, i.e. genes whose promoters are bound by Gat201, including transcriptional cofactors *GAT204* and *LIV3*. We further observed that *GAT204* and *LIV3* are also required for alkaline-repressed growth. This Gat201-dependent transcriptional pathway appears to be independent of well-studied alkaline responsive pathways: *RIM101* transcript expression does not depend on *GAT201* (Fig S7), nor does *GAT201* expression depend on *RIM101* (O’Meara et al. 2013), and our comparison of Gat201-dependent and Rim101-dependent transcriptional profiles revealed no statistical enrichment for shared targets (data not shown). Rim101 acts downstream of the cAMP pathway, and exogenous cAMP does not change the *GAT201*-dependent growth phenotypes (Fig S14). Also, the expression of genes necessary for growth in alkaline conditions, such as *PHO4* (Lev et al. 2017), *ENA1* (Idnurm et al. 2009)*, ECA1* (Ariño, Ramos, and Sychrová 2010), *CAN2*, or *CAC1* (Mogensen et al. 2006), were not dependent on *GAT201*. Although we did identify a subset of *GAT201-*dependent transcripts specific to serum within 4 hours, we did not observe an early phenotypic impact of serum on growth, suggesting that pH rather than serum is the dominant environmental signal.

The *GAT201*-dependent transcriptional profiles during alkaline-restricted growth provide some insight into the mechanisms of growth restriction. *GAT201*-dependent downregulation of ribosome biogenesis genes indicates that protein synthesis, a core pathway required for growth, is repressed downstream of *GAT201* during reactivation in RPMI. By contrast, we observed ribosomal proteins to be strongly induced in wild-type cells reactivating in rich YPD media. This shows that *Cryptococcus* grown in alkaline RPMI represses biosynthetic and replicative processes while also upregulating capsule production. In addition, others have shown that capsule synthesis is restricted to the G1 phase of the cell cycle, while budding occurs in G2 (García-Rodas et al. 2014).

We propose 3 non-exclusive hypotheses for how *GAT201* restricts growth. First, the GAT201 pathway could restrict growth directly or indirectly through transcriptional regulation of a core growth pathway. Second, the GAT201 pathway could promote capsule production, and the redirection of biosynthetic resources to capsule production could result in restricted growth. Third, some factors required for nutrient acquisition could be regulated downstream of *GAT201*, and a failure to acquire some essential nutrients would restrict growth. While nutrient acquisition could be a key contributor, we can exclude that nutrient content alone explains growth restriction, because *GAT201* and *gat201Δ* cells behave differently in identical nutrient-rich RPMI media. Modulation of media pH and buffering agents changes these growth phenotypes, and in RPMI without sodium bicarbonate at near-neutral pH *GAT201* instead promotes growth, again in media with otherwise identical nutrient composition. Alkaline conditions reduce the availability of H+ ions, which are important for the transport of nutrients across the cell membrane. We observed no *GAT201*-dependence for the expression of known H+ pumps required for alkaline growth, however several transmembrane transporters are upregulated in *GAT201* cells compared to gat201Δ. Future work will need to investigate these mechanisms.

In contrast to *C. neoformans*, ascomycete pathogens are more alkaline tolerant: *Aspergillus* species are tolerant to pH 11 (Wheeler, Hurdman, and Pitt 1991), and *Candida* species can tolerate alkaline conditions ranging from pH 10, for *C. albicans*, to pH 13 for *C. auris* and *C. parapsilosis* (Heaney et al. 2020). In ascomycetes, alkaline growth is enabled by Rim101/PacC (Selvig and Alspaugh 2011). Likewise, Rim101 is required for *C. neoformans* growth at pH 8-8.5 (Ost et al. 2015). However, *C. neoformans* also employs Rim101-independent mechanisms to mediate alkaline tolerance, for example, the sterol homeostasis pathway regulated by the transcription factor Sre1 (Brown et al. 2020). Interestingly, the basidiomycete plant pathogen *Ustilago maydis* also exhibits alkaline-restricted growth that is Rim101-independent, but the causative pathway is unknown (Cervantes-Montelongo, Aréchiga-Carvajal, and Ruiz-Herrera 2016).

Our predictions of Gat201 homologous proteins in basidiomycetes and ascomycetes also suggest hypotheses for future investigation. It would be interesting to test if the syntenic homologous protein in *Ustilago maydis*, UMAG_04076, is involved in alkaline restricted growth (Cervantes-Montelongo, Aréchiga-Carvajal, and Ruiz-Herrera 2016). The predicted *Neurospora* homologous protein, Sub-1, co-regulates genes downstream of the light responsive white collar complex, connecting light responses and fungal development (Chen et al. 2009). The *Aspergillus* homologous protein, NsdD, is a crucial regulator of sexual development (Han et al. 2001; Szewczyk and Krappmann 2010). Brg1 in *Candida albicans* is required for hyphal growth, biofilm formation and virulence (Su et al. 2018). Collectively, this suggests that Gat201 may be part of a conserved family of GATA transcription factors that regulate proliferation and morphology in response to environmental stimuli. Defining the regulatory targets, co-factors, and upstream signaling pathways leading to Gat201 family activation in different species would reveal the degree of functional conservation.

In conclusion, we have found that *GAT201* is part of an alkaline-restricted growth pathway that responds to environmental signals, including alkaline pH, to restrict cell proliferation and promote the synthesis of defensive capsule (Fig 7). The alkaline-restricted growth pathway is independent of the Rim101 pathway that promotes growth at weak alkaline pH, as well as independent of cAMP. We see over 10-fold increase in *GAT201* mRNA abundance within 30 minutes of inoculation in RPMI media, suggesting the existence of fast-acting upstream pathway components that induce *GAT201* transcription and/or stabilise the *GAT201* transcript. Downstream, Gat201 regulates hundreds of targets, including other transcription factors that are implicated in *Cryptococcus* virulence, as well as many poorly characterised genes. So far, the only other genes that we know to be required for alkaline-restricted growth encode co-factors *GAT204* and *LIV3*. Future work will map pathway components using forward genetic approaches that exploit growth conditions where the functional Gat201 pathway renders *Cryptococcus* inviable.

**Fig 7.**
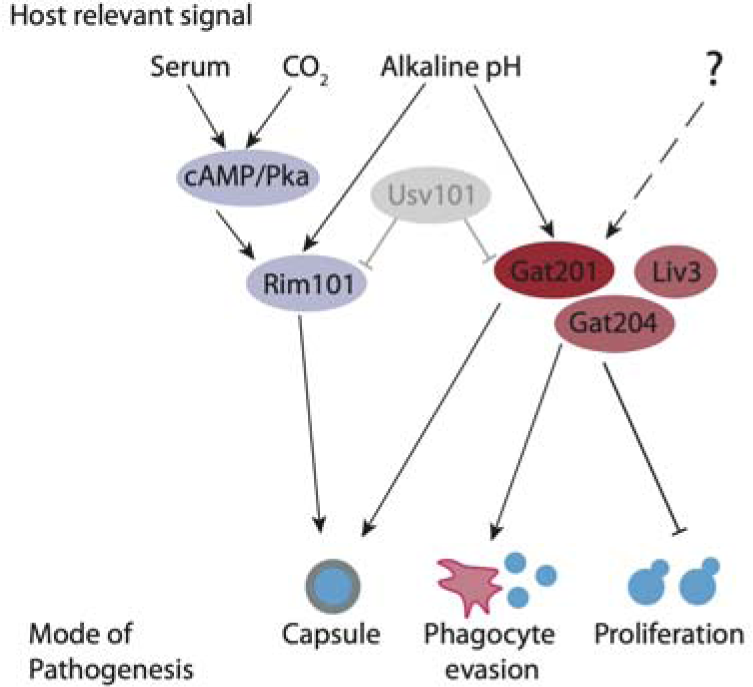
The Gat201 pathway promotes Cryptococcus virulence and represses proliferation. Gat201 acts in parallel to serum-responsive cAMP/Pka pathway and the major pH-responsive Rim101 pathway. Gat201 requires mutual activators, Gat204 and Liv3, to suppress proliferation.

## Materials and Methods

### Strains, media, and growth conditions

#### RNA-seq Dataset 1

Wild-type *C. neoformans* H99 (J. R. Perfect, Lang, and Durack 1980) yeast were gifts from Andrew Alspaugh, Duke University, NC, USA and were maintained on YPD agar plates at room temperature. For stationary phase, cells were inoculated into multiple tubes of 5 ml liquid YPD (1% yeast extract, 2% Bactopeptone, 2% Dextrose) and incubated for 5 days at 200 rpm, 30°C. On day 5, the temperature was reduced to 25°C and the cells were allowed to adjust for >4 hours. 1mL of this culture was collected for RNA extraction as the “0 minute” timepoint. Cells were counted by hemocytometer at approximately 3.5 x 10^8^ cells. Four aliquots of 3 ml of stationary phase culture were pelleted and resuspended in conditions: 25°C YPD, 37°C YPD (prewarmed), 25°C RPMI 1640 + 10% heat-inactivated fetal calf serum (HI-FCS), and 37°C RPMI + 10% HI-FCS (prewarmed), 100 ml each, so roughly a 1:30 dilution. The precise media used was RPMI 1640 with L-glutamine, sodium bicarbonate and pH indicator phenol red (Sigma R8758). Fetal calf serum (Biosera FB-1285) was heat-inactivated by incubating at 56°C for 30 minutes, then aliquoted and stored at −20°C until required. The pH was checked at time 0 and 120 minutes and confirmed to be within physiological range for the duration of the experiment (7.0 - 8.5). Cells were incubated at 60 rpm. Another stationary phase sample was inoculated into at YPD at 30°C and grown at 150 rpm for 180 min. The two biological replicates were collected on two successive days.

#### RNA-seq Dataset 2

Wild-type KN99a and KN99alpha *C. neoformans* yeast (Nielsen et al. 2003) were maintained on YPDA agar plates (1% yeast extract, 2% Bacto-peptone, 2% Dextrose, 0.002% Adenine, 2% agar) at room temperature or stored at −80°C in 15% glycerol. Gat201 mutants from the Bahn lab transcription factor disruption collection (Jung et al. 2015) and the Madhani lab *C. neoformans* deletion collection (Liu et al. 2008) were obtained from the Fungal Genetics Stock Center (Manhattan, KS, USA; https://fgsc.net). We verified the gene disruption/deletion by PCR and Sanger sequencing. For stationary phase, cells from a single colony were inoculated into 20 ml liquid YPDA (1% yeast extract, 2% Bacto-peptone, 2% Dextrose, 0.002% Adenine) and incubated for 5 days at 200 rpm, 30°C. On day 5, 3.5 ml from each sample was added to fresh media, pre-warmed RPMI or RPMI + 10% HI-FBS, to a total volume of 100 ml, and incubated at 37°C with 60 rpm shaking. Samples were collected at 30 min, 2 hours, and 4 hours. Two biological replicates were performed for each of the four strains. RPMI 1640 with L-glutamine, sodium bicarbonate and pH indicator phenol red (Sigma R8758) and heat-inactivated serum (HI-FBS; Sigma F9665, Lot# BCCB5091) was aliquoted and stored at −20°C until required.

### RNA extraction and library preparation

#### RNA-seq Dataset 1

At each time point, 15 ml of the 100 ml culture (1 ml of stationary phase culture) was collected and immediately fixed in 7 ml Methanol (32% v/v final) in a 50 mL falcon tube on dry ice. Fixed cells were pelleted, transferred to a 1.5 ml screw-top tube in 1 mL ddH2O, and pelleted again. Pellets were lyophilized and mixed with 200 µL zirconium beads and ground on a bead-beater (Biospec 112011EUR) for 5 minutes. 1400 µL of buffer RLT (Qiagen) + 1% β-mercaptoethanol was added, and the mixture vortexed. RNA was then extracted by the Qiagen RNeasy Plant Mini Kit according to the manufacturer’s protocol for filamentous fungi, proceeding from step 4. RNA quality was checked by nanodrop and agilent bioanalyzer.

#### RNA-seq Dataset 2

At each time point, 15 ml of culture (2 ml of stationary phase culture) was collected and immediately fixed in 7 ml Methanol (32% v/v final) in a 50 mL falcon tube on dry ice. Fixed cells were pelleted, transferred to a 1.5 ml screw-top tube in 1 mL ddH2O, and pelleted again. Pellets were lyophilized, then mixed with 200 µL zirconium beads + 1 ml TRI reagent (Invitrogen) and incubated at room temperature for 5 mins followed by flash freezing in a dry ice/ethanol bath. Samples were thawed at room temperature and subjected to mechanical lysis by bead beating (using a PreCellys machine) for 3 x 10 seconds (6000 rpm) followed by a pause of 20 seconds and placed on ice for 1 minute. This was repeated 10 times. After mechanical disruption the zirconium beads were pelleted and the supernatant transferred to a QIA shredder spin column from the Qiagen RNeasy Plant Mini Kit (Qiagen, Valencia, CA, USA) and RNA was then extracted according to the manufacturer’s protocol, proceeding from step 4. Total RNA quantity and quality were assessed using nanodrop and the Fragment Analyser Automated Capillary Electrophoresis System (Agilent Technologies Inc, #5300) and the Standard Sensitivity RNA Analysis Kit, 15nt (#DNF-471).

For the experiments in Fig 1 (Dataset 1), 2 µg of RNA from each sample was used as input for RNA sequencing by the RNATagSeq protocol (Shishkin et al. 2015), with minor modifications including ribosomal RNA depleted using the Yeast RiboZero Gold kit (Illumina; now discontinued), and a random barcode added to the 2nd ligation primer, with 12 cycles of PCR. Libraries were sequenced on a Nextseq500 (Illumina).

For the experiments in Figs 3-4 (Dataset 2) 500 ng of RNA from each sample was used as input for cDNA library preparation using the QuantSeq FWD 3’ mRNA-Seq Library Prep Kit for Illumina platforms according to the manufacturer’s instructions. We spiked in 10 ng of *Saccharomyces cerevisiae* total RNA as a loading control, but did not use this spike-in information in the data analysis presented here. The QuantSeq protocol generates only one fragment per transcript, close to the 3’ end of the transcripts. cDNA fragments of ∼300 bp were purified from each library and confirmed for quality by the Fragment Analyser Automated Capillary Electrophoresis System (Agilent Technologies Inc, #5300) and the Standard Sensitivity NGS 1-6000bp Kit (#DNF-473-33). Single read sequencing was performed using the NextSeq 500/550 High-Output v2.5 (75 cycles) Kit (#20024906) on the NextSeq 550 platform (Illumina Inc, #SY-415-1002). Libraries were combined in a single equimolar pool of 56 based on Qubit and Bioanalyser assay results and run across a High-Output v2.5 Flow Cell.

RNA-sequencing data are available on GEO under accession numbers GSE133067 (Dataset 1) and GSE217345 (Dataset 2).

### Bioinformatic and statistical data analyses of RNA-seq data

Complete analysis code for the RNA-seq datasets from raw reads onwards is found on github, https://github.com/ewallace/CryptoWakeupRNASeq (Dataset 1) https://github.com/ewallace/CryptoGat201RNASeq (Dataset 2).

In summary, basic assessments of sequence data quality were performed using FastQC (Andrews and Others 2010) and MultiQC (Ewels et al. 2016). Raw sequencing reads were trimmed and filtered using cutAdapt (Martin 2011). Sequenced reads were aligned to the *C. neoformans* H99 reference sequence CNA3 (Janbon et al. 2014) using Hisat2 (Daehwan Kim et al. 2019). We used featureCounts (Liao, Smyth, and Shi 2014) to assign mapped reads per gene using the longest transcript per gene annotation from (Wallace et al. 2020). Gene expression was normalized using the regularized logarithm (rlog) function from DESeq2 (Love, Huber, and Anders 2014). We evaluated the gene expression differences using a test based on a negative binomial distribution, also in DESeq2 (Love, Huber, and Anders 2014), using a 5% false discovery rate calculated by the ‘p.adjust’ function in R using the Benjamin and Hochberg method (Haynes 2013).

Gene clustering was performed by using (1 - correlation) as a distance metric, then hierarchical clustering in UPGMA using R’s hclust function (Holmes and Huber 2019). After a lengthy iterative process in which many methods were evaluated, representative clusters of genes were selected from the hierarchical clusters using R’s cutree function with user-defined numbers of groups (k option). Full details and analysis code are in the repositories.

Differentially expressed genes and gene clusters were subjected to GO term enrichment analysis using the online resources at FungiDB (Basenko et al. 2018).

### Strain Construction

To complement gat201Δ, theGAT201 gene including native promoter, terminator, and introns with a HYG selection marker (pGAT201-cGAT201-tGAT201-HYG) was integrated into a genomic safe haven locus 4 on chromosome 7 (Erpf, Stephenson, and Fraser 2019) in the Madhani lab *gat201Δ* strain. We made the integration constructs using modular cloning by Möbius assembly (Andreou and Nakayama 2018), the details of which we will explain in another publication. A full plasmid map is included in supplementary table S4. We integrated this construct using a *Cryptococcus* CRISPR-Cas9 system (Huang et al. 2022).

### Microscopy

For stationary phase, cells were revived from −80°C glycerol stocks on YPD agar and within two days single colonies were inoculated into 10 ml liquid YPD (1% yeast extract, 2% Bactopeptone, 2% Dextrose) and incubated for 5 days at 200 rpm, 30°C. On day 5, the temperature was reduced to 25°C and the cells were allowed to adjust for >4 hours. Cells were counted by hemocytometer and 1 ml of the pellet (10^6^ cells) was collected, washed 1× with PBS, and split into four tubes, then resuspended in the appropriate pre-warmed medium as indicated to a final volume of 10 ml each. Cells were incubated in the indicated condition for 120 minutes, and then the entire pellet was collected and fixed with 4% methanol free formaldehyde (Pierce) for 10 minutes, then washed 3x with PBS. India ink (Remel) slides were prepared and cells were imaged using an inverted Zeiss AxioObserver Z1 with a Plan-Neofluor 40X/1.3Lnumerical aperture (NA) oil immersion lens objective (Carl Zeiss) and a 16-bit CoolSNAP H2 charge-coupled-device (CCD) camera (Photometrics). For each figure, three biological replicates were initiated using independent stationary cultures and collected serially on the same day. The entire experiment was performed independently twice.

### Growth Curves

Cells from a single colony were inoculated into 5 ml liquid YPDA (1% yeast extract, 2% Bacto-peptone, 2% Dextrose, 0.002% Adenine) and incubated for 20 hours at 180 rpm, 30°C. Cells were washed with ddH2O and resuspended in the required volume of the appropriate media to an initial OD 595 nm of 0.2. Wells in the microplate were filled with this suspension (200 µl in each well). The absorbance in each well was measured at 595 nm at given intervals (10 minutes) with shaking (300 rpm for 1 minute) directly prior to reading. Reference measurements were performed on the outer wells where 200 µl of media only was added. The microplate was Incubated in Tecan Infinite® 200 Pro plate reader at 37°C for 48 or 72 hours. Cells were grown as stated in RPMI 1640 (Sigma R8758) with or without heat-inactivated serum (Sigma F9665), YPDA, for figures except where noted. For Fig S12, we used CO_2_-independent media, buffered with mono and dibasic sodium phosphate and β-glycerophosphate (Gibco™ / ThermoFisher 18045088). For Figs 4, S13, and S14, we used RPMI media without phenol red and without NaHCO_3_ (Sigma R7855), adding dibutyryl cAMP (Sigma D0627) or NaHCO_3_ from aqueous stock solutions to the indicated final concentration. For these we were particularly careful to rapidly prepare media and then inoculate cells for growth curves, for reproducible pH of the growth media.

### Colony forming unit assay

Colony forming unit assay is an *in vitro* cell survival assay based on the ability of a single cell to grow into a colony. Cells from a single colony were inoculated into 25 ml liquid YPDA (1% yeast extract, 2% Bacto-peptone, 2% Dextrose, 0.002% Adenine) and incubated for 5 days at 150 rpm, 30°C. Cells were washed with ddH2O and resuspended in a total volume of 20 ml RPMI at an OD 595 nm of 0.1 for each strain. Cultures were incubated at 37°C, 60rpm and samples were collected at 12 hour intervals (0-60 hours). Serial dilutions were prepared for each sample collected from each strain down to 10^−4^, 100 μl of dilutions 10^−3^ and 10^−4^ were plated onto YPDA agar plates and incubated at 30°C for 48 hours. Plates were imaged using an ImageQuant 800 (Amersham/Cytiva, settings: Colorimetric, OD measurement, Auto exposure, Capture area = 160 x 220 nm) and the resulting colonies were counted. Three biological replicates were collected for each strain.

### RT-qPCR

Cells from a single colony were inoculated into 5 ml liquid YPDA (1% yeast extract, 2% Bacto-peptone, 2% Dextrose, 0.002% Adenine) and incubated for 20 hours at 180 rpm, 30°C. Cells were washed with ddH2O and resuspended in 100 ml RPMI (Sigma R8758) at an initial OD 595 nm of 0.1, incubated at 37°C, 150 rpm for 7 hours. Cells were fixed in methanol and RNA extracted using mechanical disruption in TRizol with zirconium beads followed by the Qiagen Plant and Fungal Extraction Kit. 100 ng of purified RNA was used from each sample to synthesize cDNA using Superscript IV Reverse Transcriptase (Invitrogen) and random primers (NEB). Samples were DNase treated prior to reverse transcription. QPCR was carried out using Brilliant III Ultra fast SYBR Green qPCR mix (Agilent) with appropriate target gene primers. mRNA expression of GAT201 (Forward primer: 5’-ACCACGAGTCTTGGGATAGA-3’, Reverse primer: 5’-CTGGGTGTTCGGGATAAAGTAG-3’), GAT204 (Forward primer: 5’-CCACCTCTTCCTTCCTTGTTAAA-3’, Reverse primer: 5’-GTCTGCCATCGTCGTACTAATG-3’), and LIV3 (Forward primer: 5’-CCTCTTCCACTTCCACATCAA-3’, Reverse primer: 5’-GGTCTCGGCACAGCATATT-3’). Test genes were compared to 3 reference genes ACT1 (Forward primer: 5’-GTGGTTCTATCCTTGCCTCTTT-3’, Reverse primer: 5’-CACTTTCGGTGGACGATTGA-3’), GPD1 (Forward primer: 5’-TCGAGCAACGTCTTGGTATC-3’, Reverse primer: 5’-GCTCTCCATCCTCCTTGTTT-3’) and SRP14 (Forward primer: 5’--3’, Reverse primer: 5’--3’). The data were analysed with tidyqpcr (Wallace and Haynes 2022).

### Data analysis

Data analysis scripts and raw data for budding index assay, growth curve assay, CFU assay, and RT-qPCR are in the repository https://github.com/ewallace/CryptoGat201_2023_suppdata. Scripts and raw data for the homology analysis are in the repository https://github.com/ewallace/Gat201homology_2022/. Data were analysed in the statistical open-source language R (R Core Team 2023), making extensive use of the tidyverse for data manipulation (Wickham et al. 2019) and ggplot2 for figures (Wickham 2009). Additional figures were prepared in Inkscape (The Inkscape Team, https://inkscape.org/).

## Acknowledgements

E.S.H. was supported by a Fellowship from the Daphne Jackson Memorial Trust, funded by the University of Edinburgh and the Biotechnology & Biological Sciences Research Council. E.R.B. and E.W.J.W. are each supported by Sir Henry Dale Fellowships jointly funded by the Wellcome Trust and the Royal Society (211241/Z/18/Z to E.R.B., 208779/Z/17/Z to E.W.J.W.). Collection of Dataset 1 was funded by a Research Incentive Grant from the Carnegie Trust for the Universities of Scotland to E.W.J.W., and by the European Union’s Horizon 2020 research and innovation programme under Marie Skłodowska-Curie grant agreement no. 661179 to E.W.J.W. We acknowledge funding from the MRC Centre for Medical Mycology at the University of Exeter (MR/N006364/2 and MR/V033417/1), and the NIHR Exeter Biomedical Research Centre. Additional work may have been undertaken by the University of Exeter Biological Services Unit. The views expressed are those of the author(s) and not necessarily those of the NIHR or the Department of Health and Social Care. We thank Hiten Madhani for gifts of deletion strains, supported by NIH grant (R01AI100272). We thank Andrew Cassidy for sequencing RNA-seq dataset 1 at the Tayside Centre for Genomic Analysis, University of Dundee. We thank Richard Clarke, Angie Fawkes, and Lee Murphy, for sequencing RNA-seq dataset 2 at the Genetics Core of the Edinburgh/Wellcome Trust Clinical Research Facility. We thank the Wallace lab and Ballou lab for comments and feedback on the project over the years.

## Notes

https://www.ncbi.nlm.nih.gov/geo/query/acc.cgi?acc=GSE133067

https://www.ncbi.nlm.nih.gov/geo/query/acc.cgi?acc=GSE217345

https://github.com/ewallace/CryptoWakeupRNASeq

https://github.com/ewallace/CryptoGat201RNASeq

https://github.com/ewallace/CryptoGat201_2023_suppdata

https://github.com/ewallace/Gat201homology_2022/

### Competing Interest Statement

The authors have declared no competing interest.

### Summary of Updates

- New data added on dependence of growth on pH / NaHCO3, on cyclic AMP pathway, and genetic interactions between GAT204 and LIV3 - New phylogenetic analysis added of GAT201. - Several clarifications to text. - These revisions were made in response to a journal review process.

