## Supplementary Figures S1-S15 for "A trade-off between proliferation and defense in the fungal pathogen *Cryptococcus* at alkaline pH is controlled by the transcription factor GAT201"

Elizabeth S. Hughes^1^, Laura R. Tuck^1^, Zhenzhen He^1^, Elizabeth R. Ballou^2^, Edward W.J. Wallace^1^.

1. Institute for Cell Biology, and Centre for Engineering Biology, School of Biological Sciences, The University of Edinburgh. 2. MRC Centre for Medical Mycology, The University of Exeter.

Contains Supplementary Figs S1-S15.

**Fig S1, related to Fig 1C: Principal Component Analysis of RNA-seq dataset 1.** Note PC 1 vs 2 panel is a repeat of 1C.

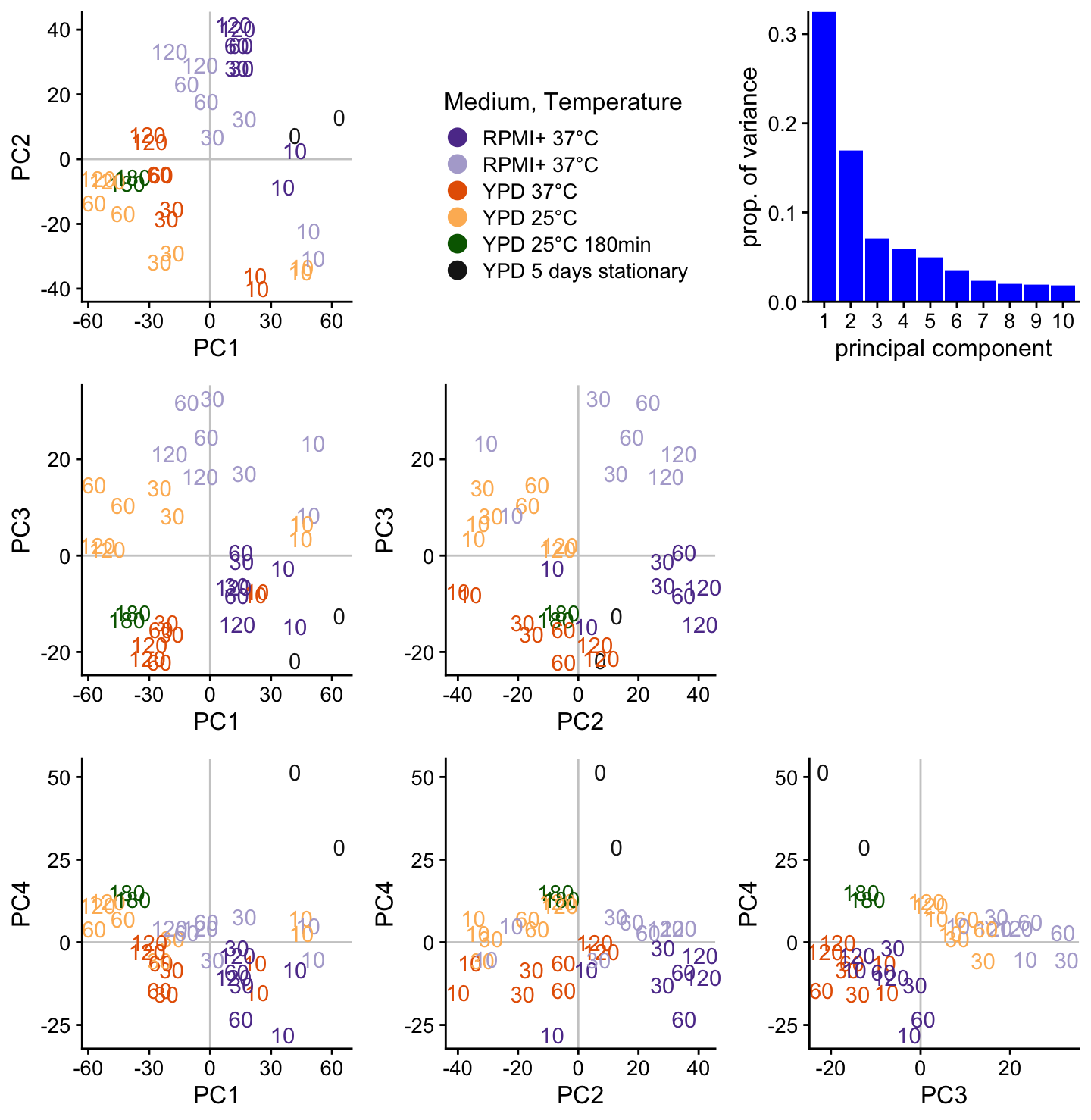

**Fig S2, related to Fig 1E: Thirty representative genes show distinct expression patterns, again in log2 fold-change per gene.** See Fig 1 legend for details of the experiment.

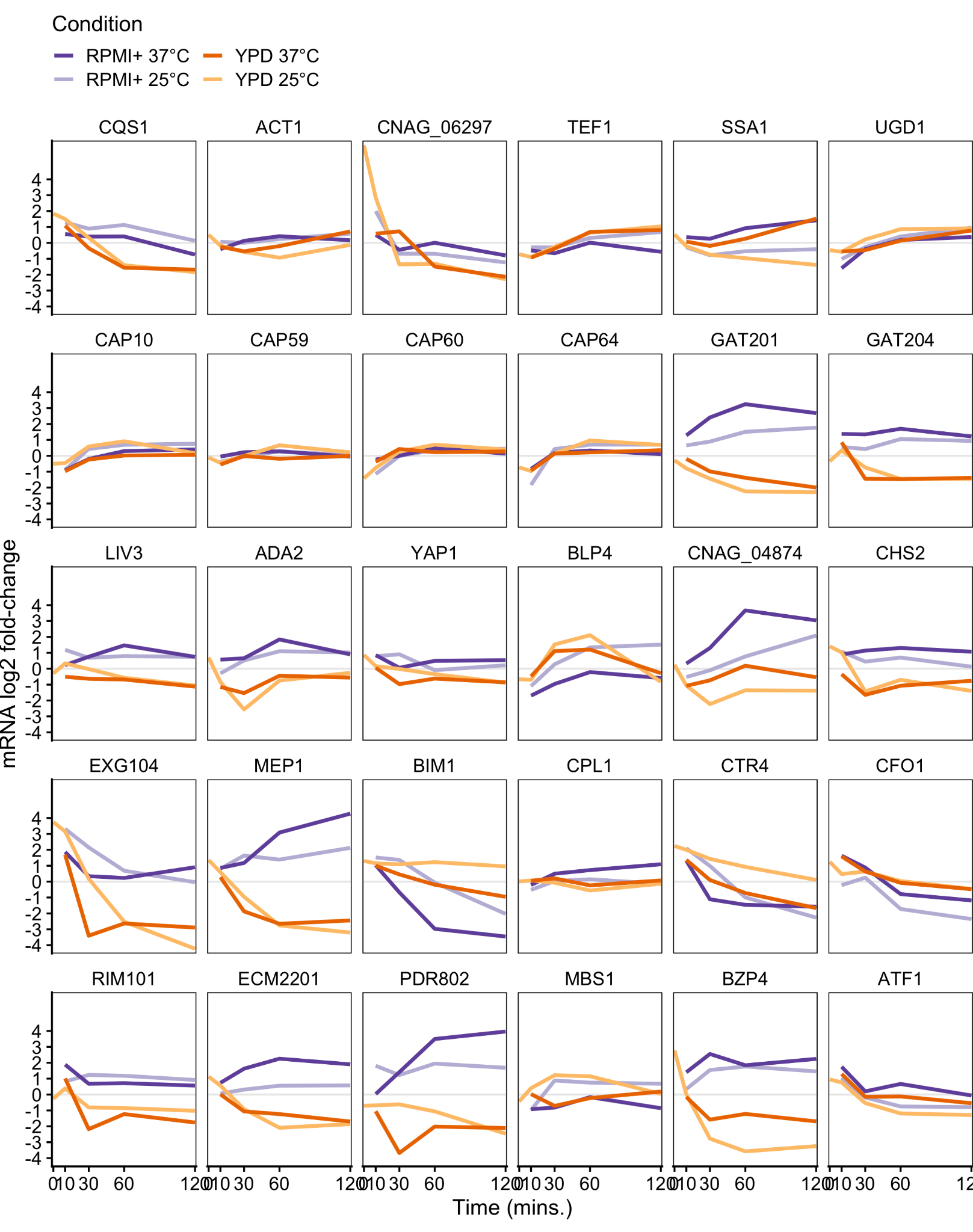

**Fig S3, related to Fig 2: GAT201 promotes capsule biosynthesis and represses budding in RPMI medium (without serum) at 37°C 2 hours after inoculation.** There is no obvious difference in phenotype in rich YPD medium. Micrographs show GAT201 (H99), gat201∆m, and complemented GAT201-C1 strains, stained with India Ink. Scale bar is 10µm.

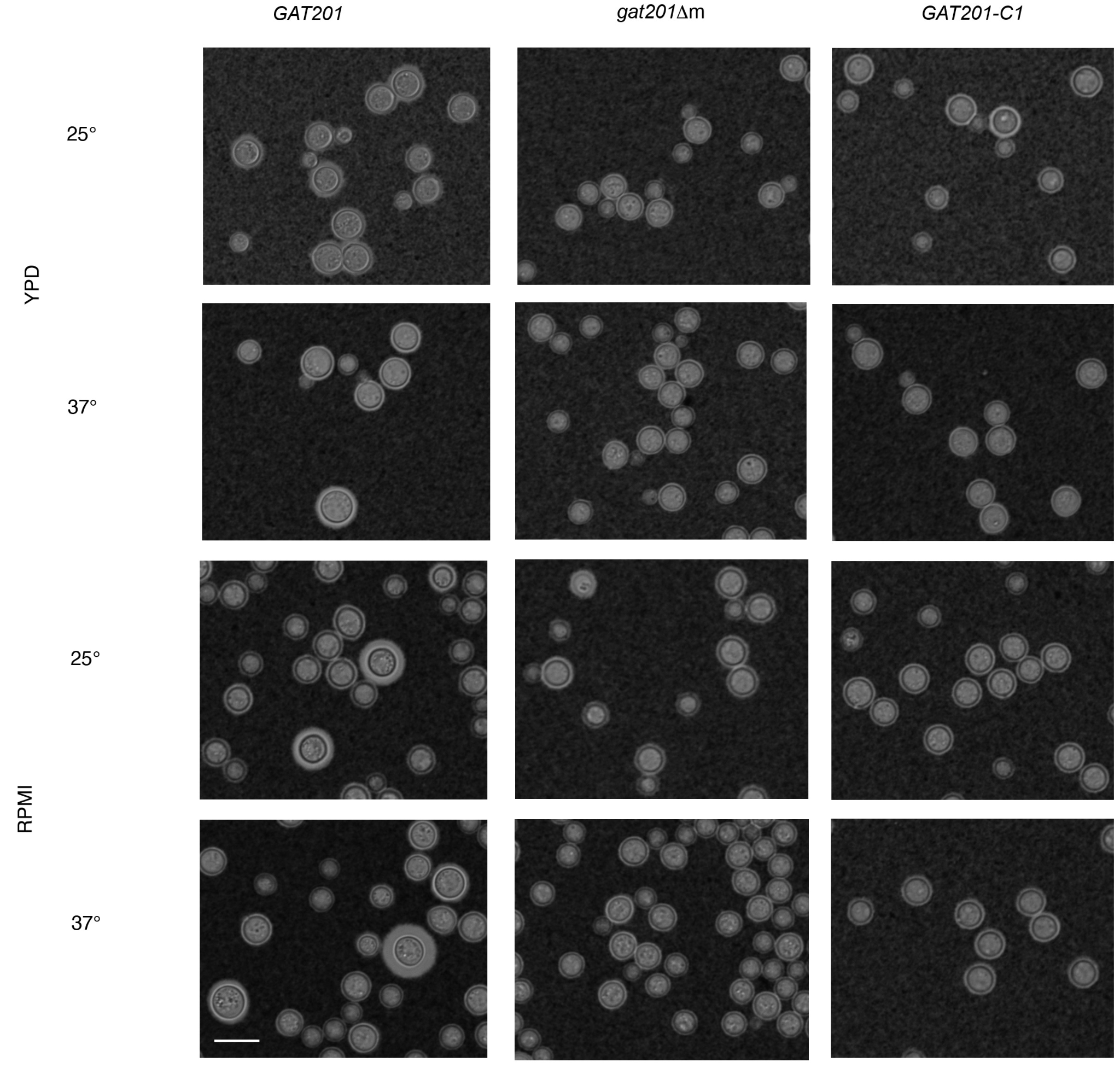

**Fig S4, related to Fig 2: Partial complementation of Gat201 mRNA expression shown by RT-qPCR of GAT201, GAT204, and LIV3 genes compared to 3 reference genes (ACT1, GPD1, SRP14).** In complemented strains, GAT201 mRNA abundance is roughly 10x lower (log2 fold-change ~3.3) than in wild-type *GAT201* (KN99alpha). Cultures were inoculated from overnight growth in YPD and grown in RPMI at 37°C for 7 hours. The figure shows log2 fold-change (∆∆Cq) values from 3 biological replicates (median of 3 technical replicates), and a mean value across the biological replicates. The low value of GAT201 detected in *gat201∆m* represents background, and was only detected at all in 2 out of 3 biological replicates.

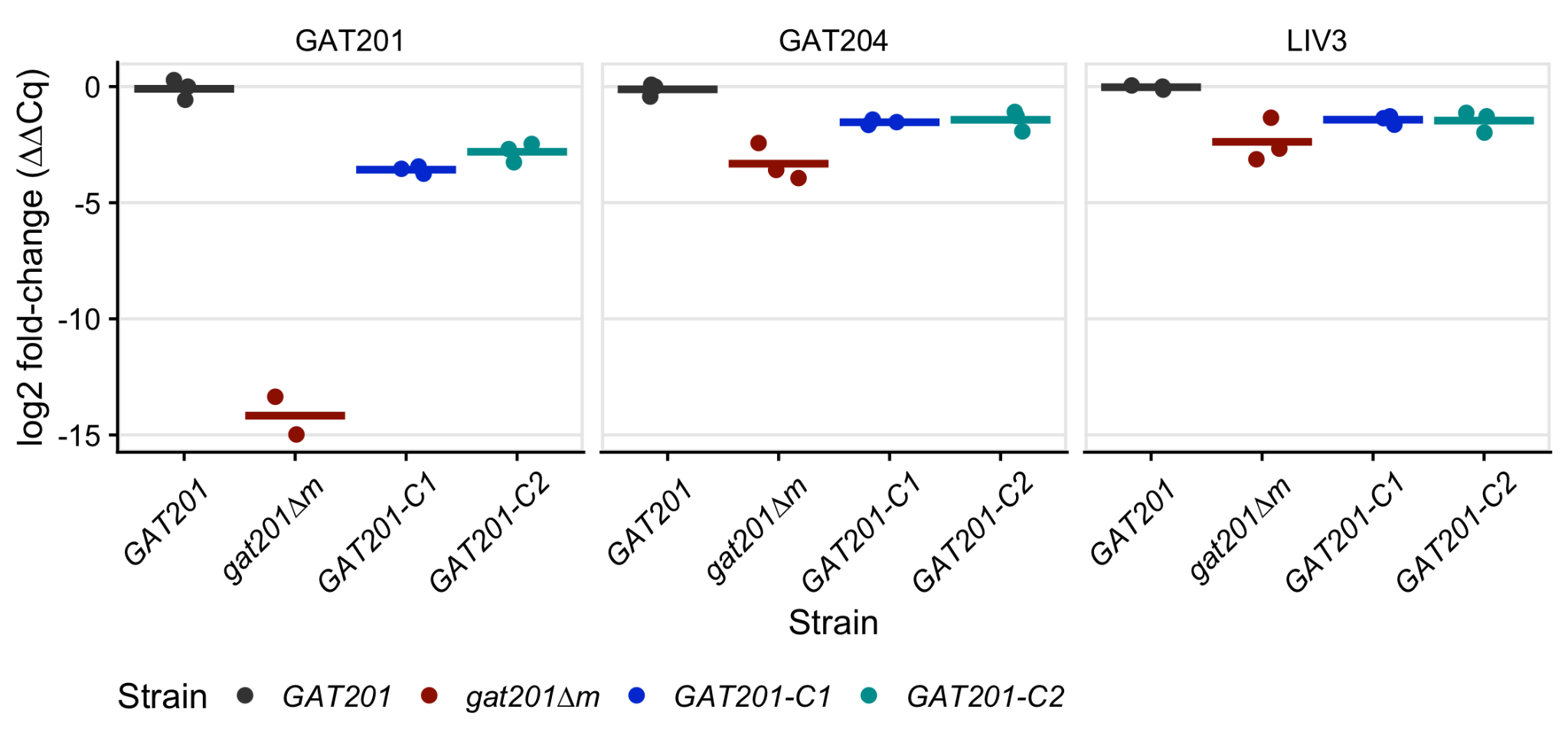

**Fig S5, related to Fig 3A: *GAT201* promotes capsule biosynthesis and represses budding in RPMI medium both with and without serum at 37°C, 2 hours after inoculation.** Strains are *GAT201* (KN99alpha) and *gat201∆m*, here a wider field of view is shown than in Fig 3A.

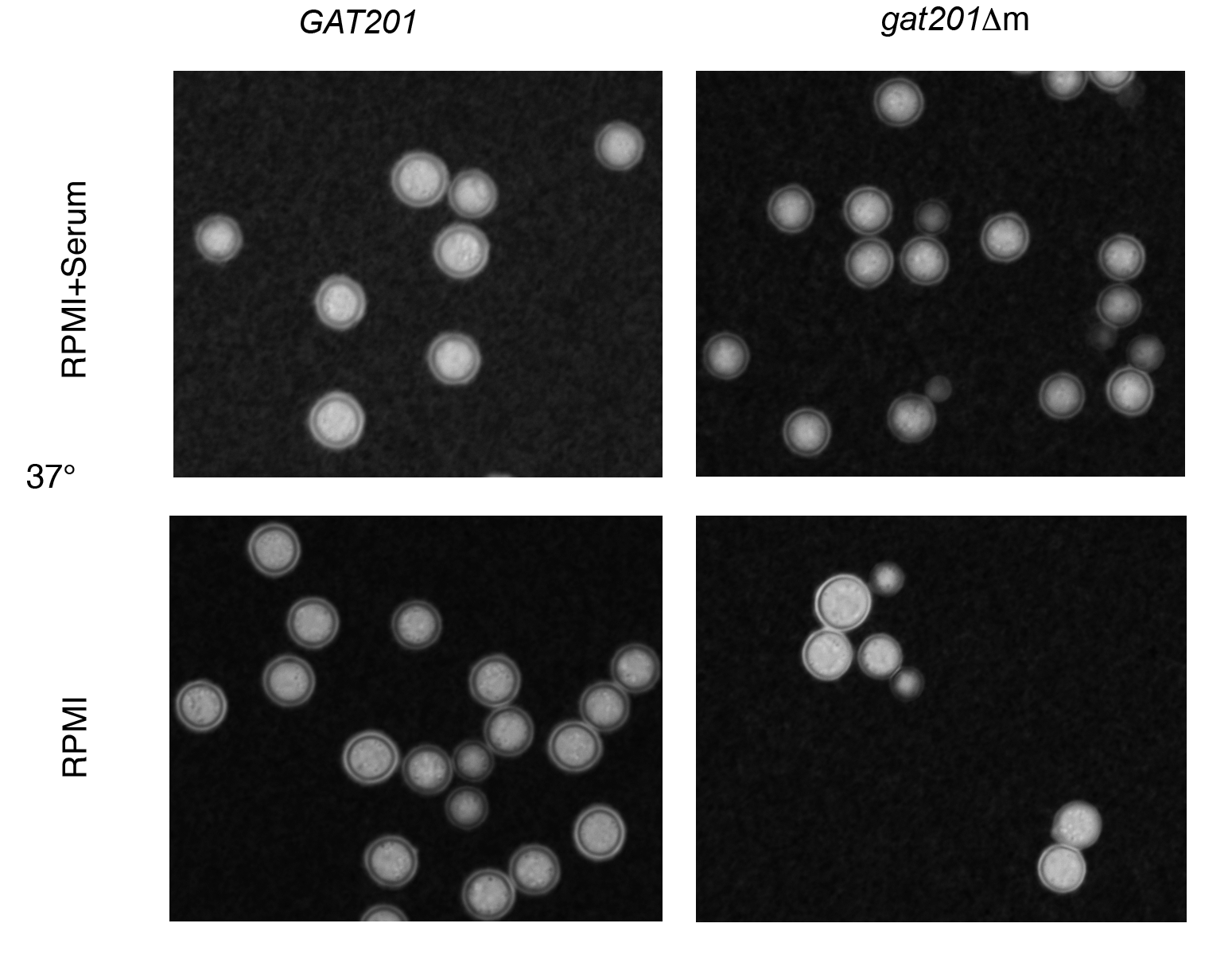

**Fig S6, related to Fig 3B: Principal Component Analysis of RNA-seq dataset 2, comparing *GAT201* (wild-type) to *gat201∆* strains.** Note PC 1 vs 2 panel is a repeat of 3B.

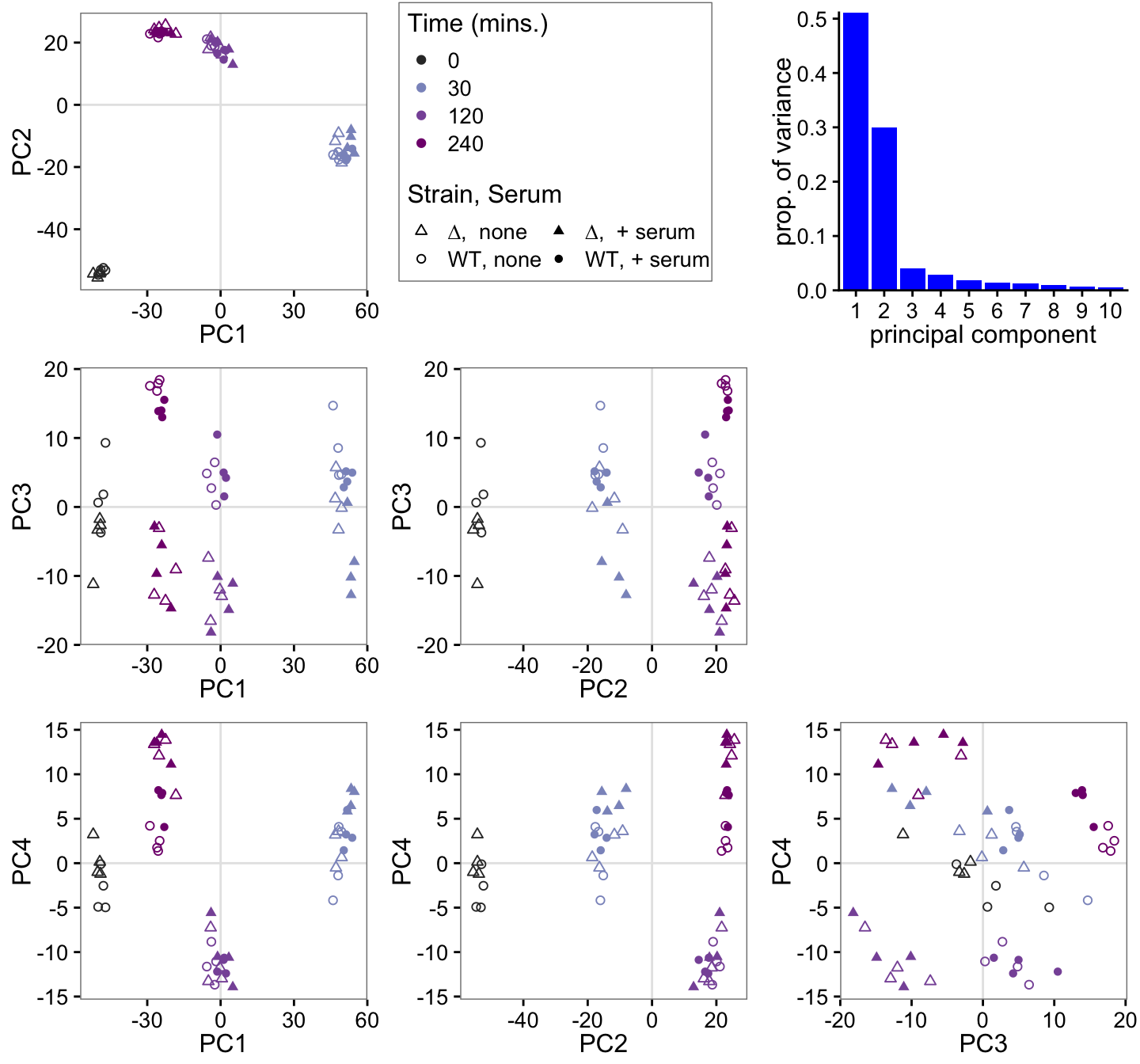

**Fig S7, related to Fig 3D: Thirty representative genes show distinct expression patterns and GAT201 dependence, again in log2 fold-change per gene.** See Fig 3 legend for details.

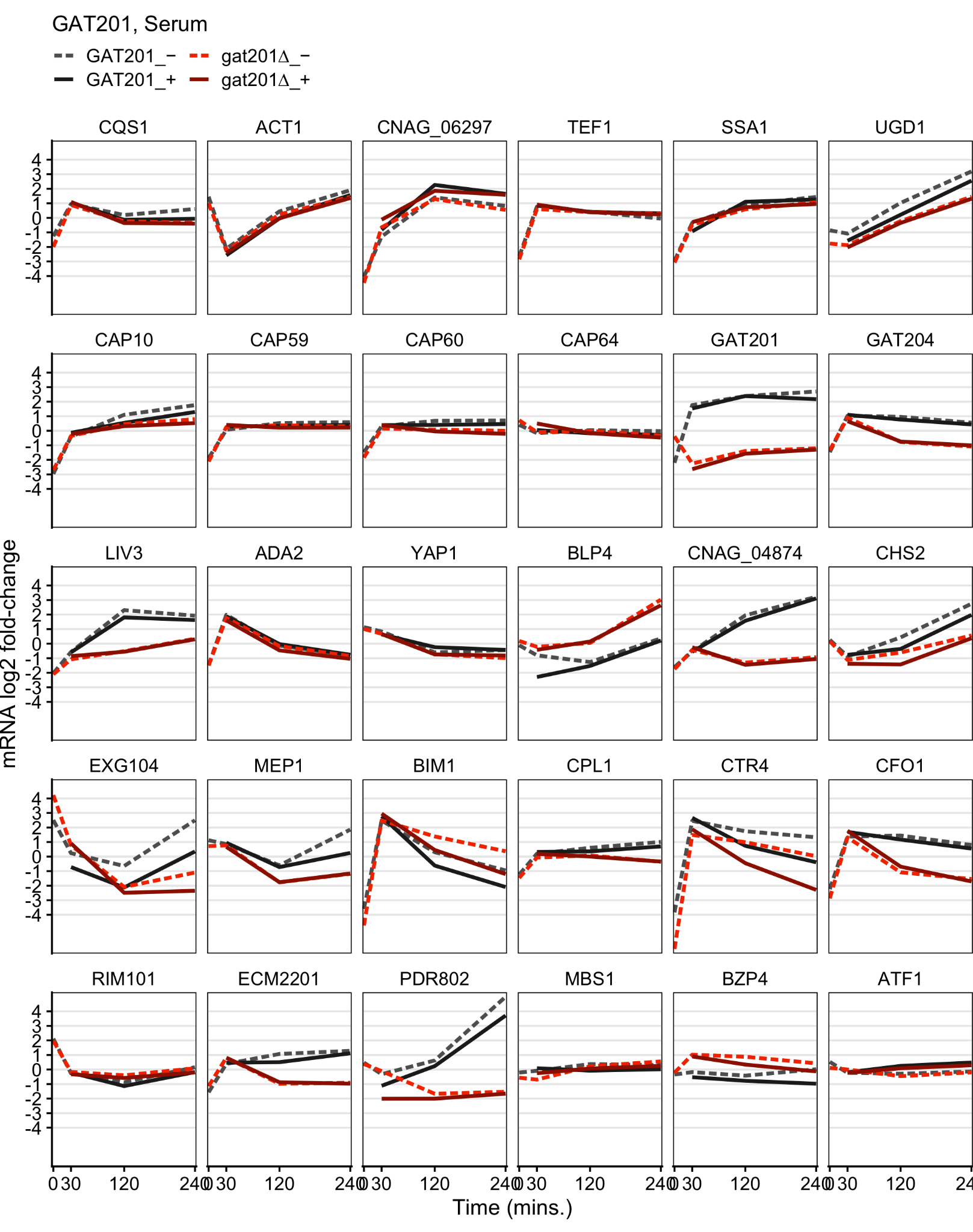

**Fig S8, related to Fig 3. Serum affects expression of a small set of transcripts.** A, Number of differentially expressed genes (2x changed, 5% FDR) dependent on serum at each timepoint in *GAT201* and *gat201∆* strains. B, Serum-dependent log2 fold-change and p-values in *GAT201* at 4 hours. C, Serum-dependent log2 fold-change and p-values in *gat201∆* at 4 hours.

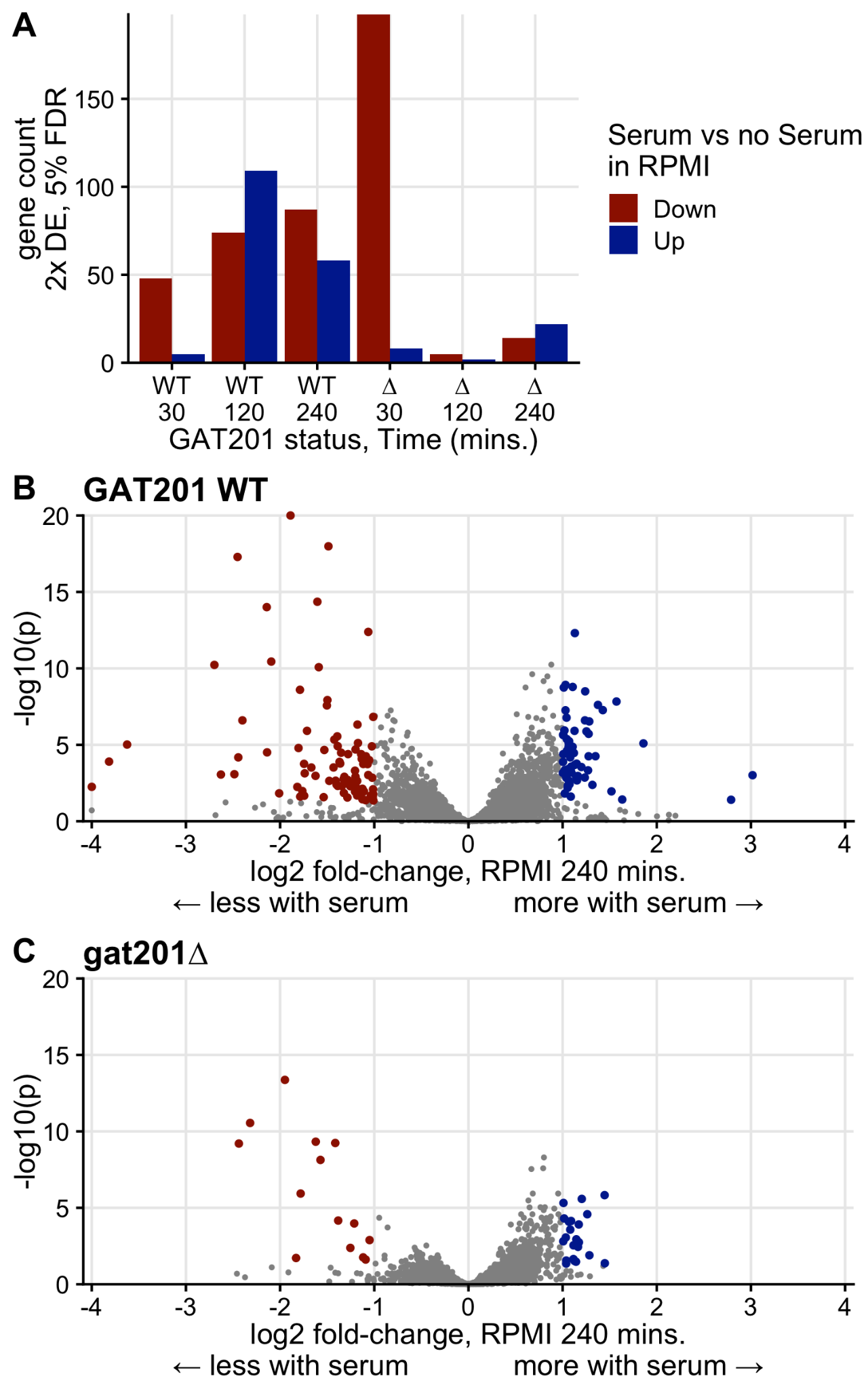

**Fig S9: Comparison of differential expression in dataset 1 (Wakeup) and dataset 2 (Gat201), differential expression between 0 hours (pre-inoculation timepoint) and 2 hours in RPMI + serum for wild-type strain.** The lowest 10% expressed transcripts in both of these datasets (by baseMean from DESeq2) were filtered out. 30 genes shown in Figs S2, S7 and discussed in the text are highlighted.

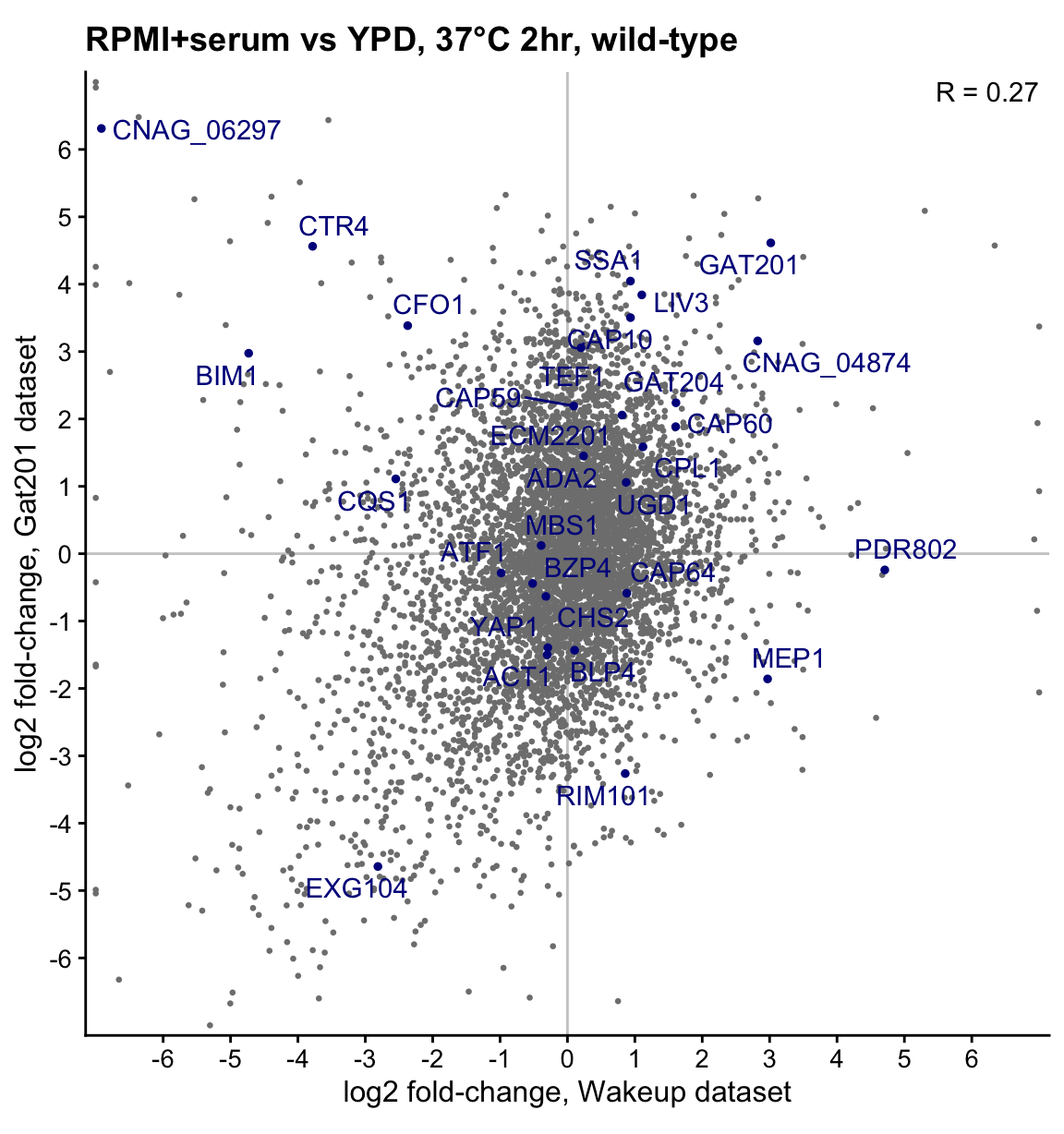

**Fig S10, related to Fig 4: Growth curves of mutants in the Gat201 pathway.** Panels A,B,C show growth curves, all 4 biological replicates in 3 technical replicates, of OD595 corrected for values of blank wells. Individual wells are plotted as faint lines, and smoothing spline for each strain plotted as thick lines. The summaries from panel A are also shown in Fig 4E.

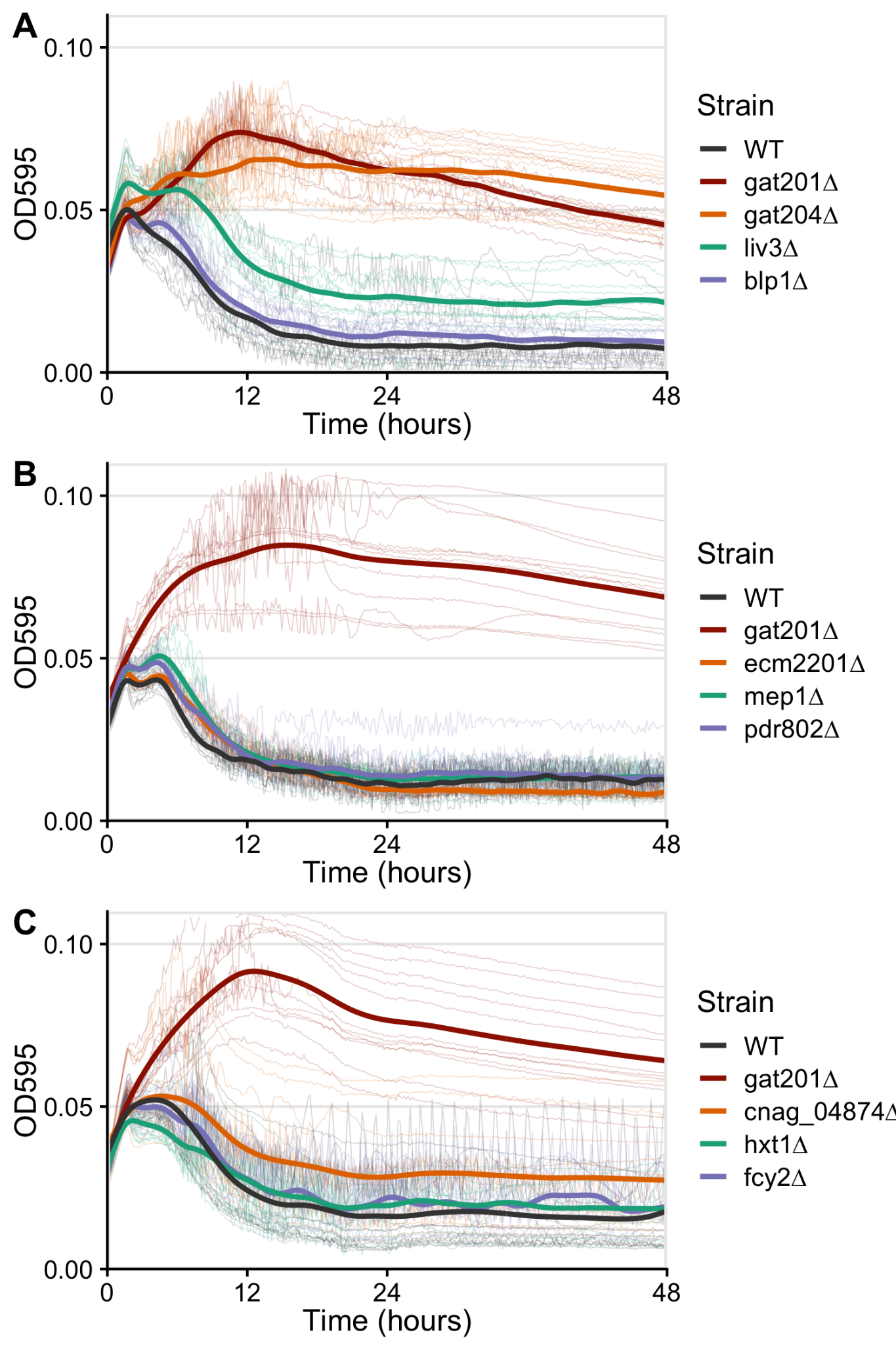

**Fig S11, related to Fig 4: The double mutant gat201∆ liv3∆ has similar phenotypes to single mutants of partially permitting growth in RPMI.** Panel shows growth curves, all 3 biological replicates in 3 technical replicates, of OD595 corrected for values of blank wells. Individual wells are plotted as faint lines, and smoothing spline for each strain plotted as thick lines.

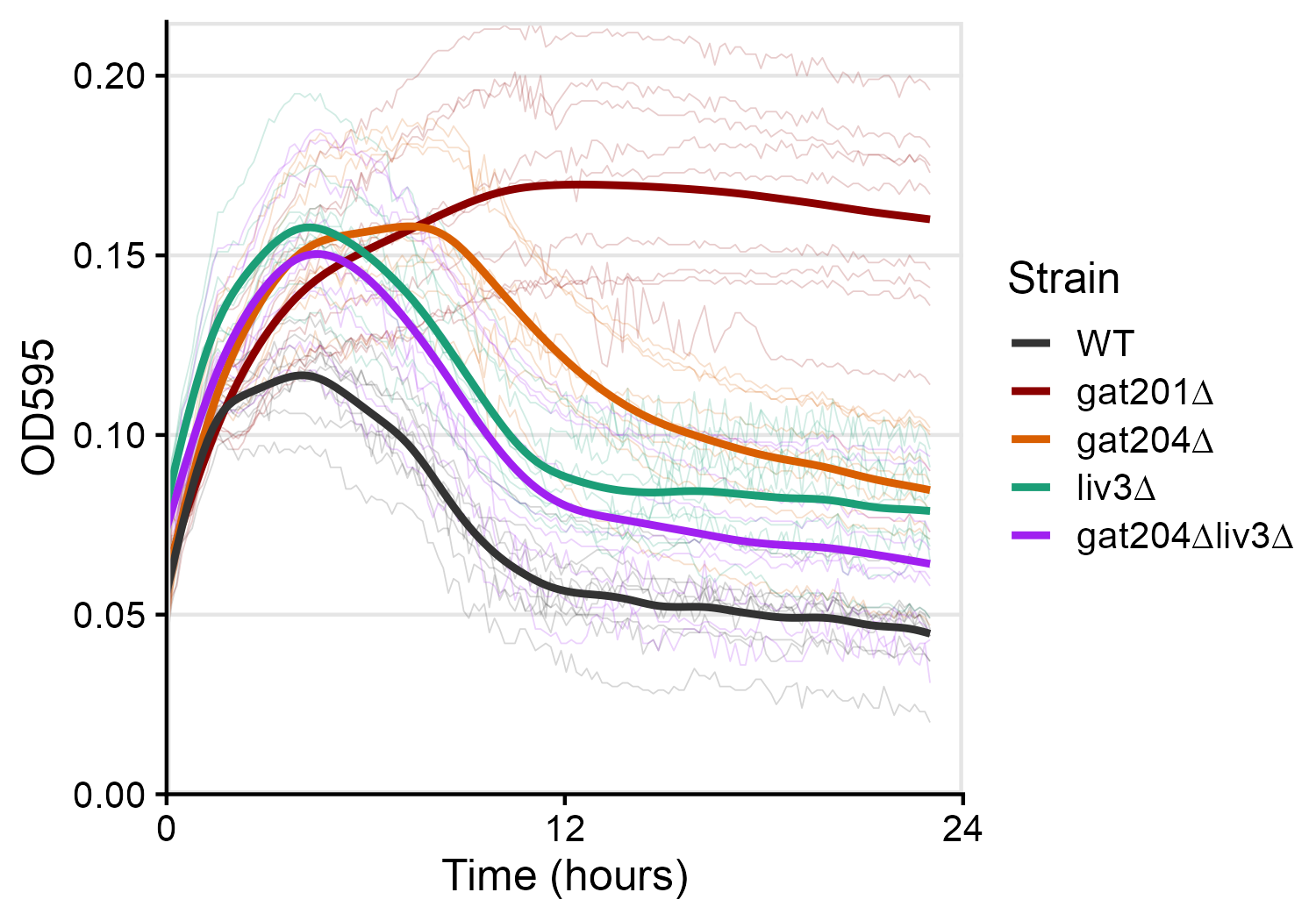

**Fig S12: GAT201 has a minor effect on growth in media buffered to near-neutral pH with mono and dibasic sodium phosphate and β-glycerophosphate.** Cells were grown as in Fig 2C and their OD595 measured, but in Gibco™ CO_2_-independent media. Individual wells are plotted as faint lines, and smoothing spline for each strain plotted as thick lines.

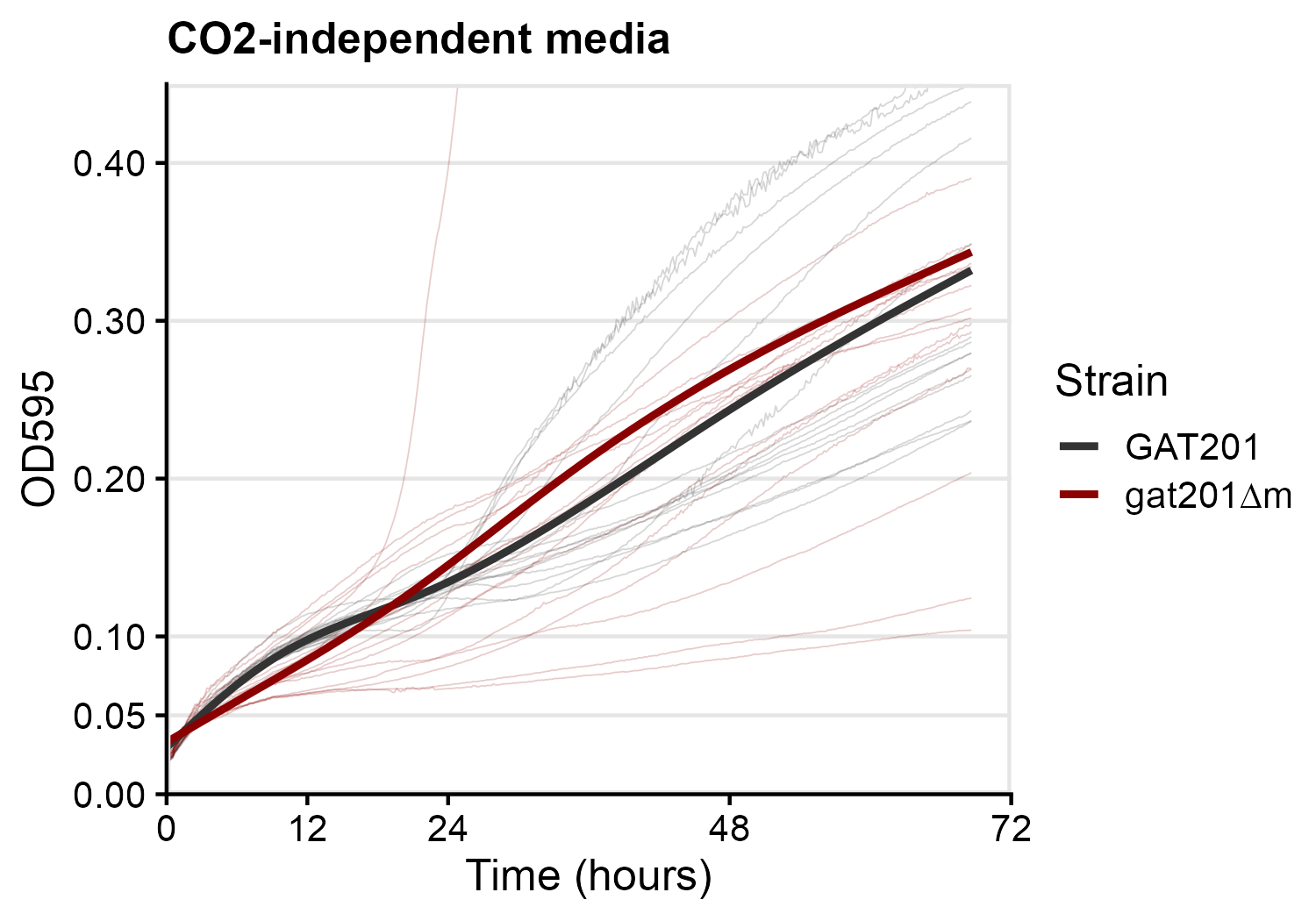

**Fig S13, related to Fig 5: The effect of *GAT201* on growth depends on sodium bicarbonate (NaHCO_3_)**. Panel shows growth curves, each column showing a single biological replicate (Biol. Rep.), and within those 3 technical replicates for each strain, of OD595 corrected for values of blank wells. Individual wells are plotted as faint lines, and smoothing spline for each strain plotted as thick lines. The summary splines for biological replicate 3 are also shown in main Fig 5.

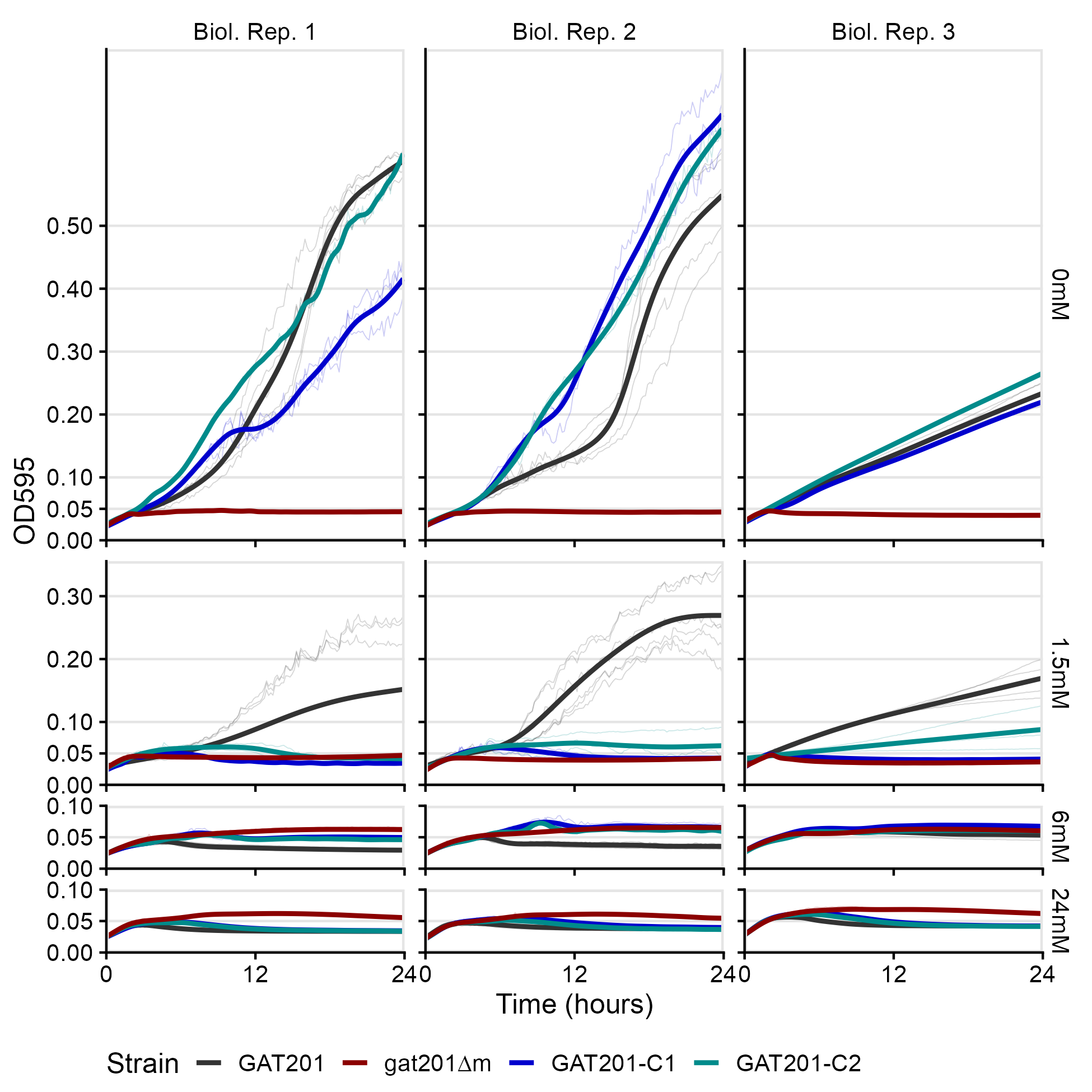

**Fig S14: Growth phenotypes due to sodium bicarbonate (NaHCO_3_) and GAT201 are not dependent on cyclic AMP (cAMP).** Fig shows growth curves of wild-type *GAT201* and deletion mutant *gat201∆m*, grown in RPMI. with no added NaHCO_3_ (top row, “−”) or 24mM NaHCO_3_ (“+”), and with either mock H_2_O addition, exogenous cAMP (N6-2'O-dibutyryl-cAMP; dbcAMP) to stimulate the pathway, or sodium butyrate (Nabut) as an additional control.

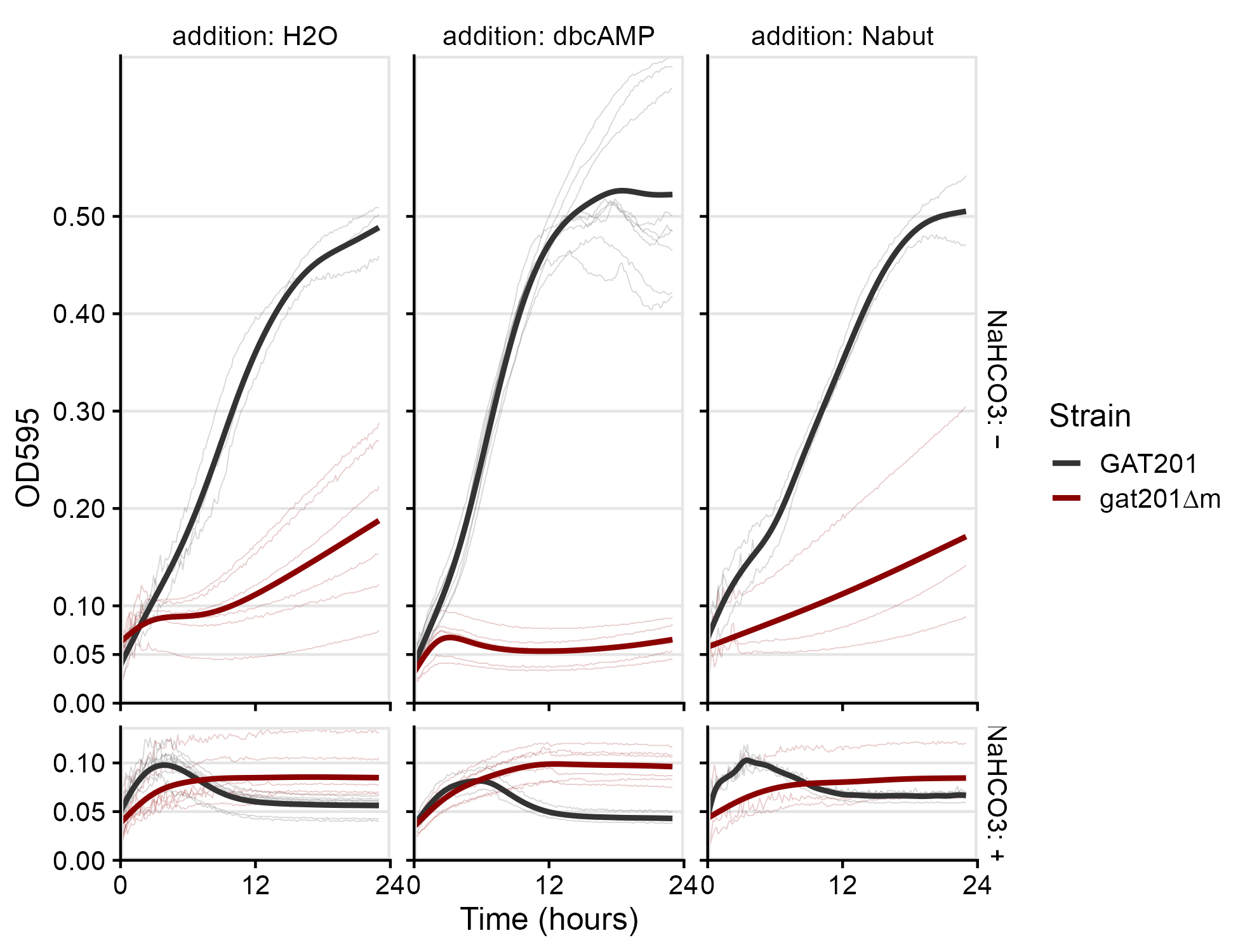

**Fig S15: Extended phylogenetic analysis of Gat201 and related GATA-domain zinc finger proteins shows that fungal *C. neoformans* Gat201, *C. albicans* Brg1, *A. nidulans* NsdD, N. crassa *sub-1*, and amoebal *D. discoideum* gtaI, form a subfamily.** A, Phylogenetic tree calculated by IQ-TREE from a full-length protein alignment by MAFFT, highlighting the groups of *C. neoformans* Gat201-like, *S. cerevisiae* Gat2-like, and *C. neoformans* Gat204-like proteins. B, Multiple sequence alignment by MAFFT of the GATA zinc finger domain from the same proteins in the same order as panel A.

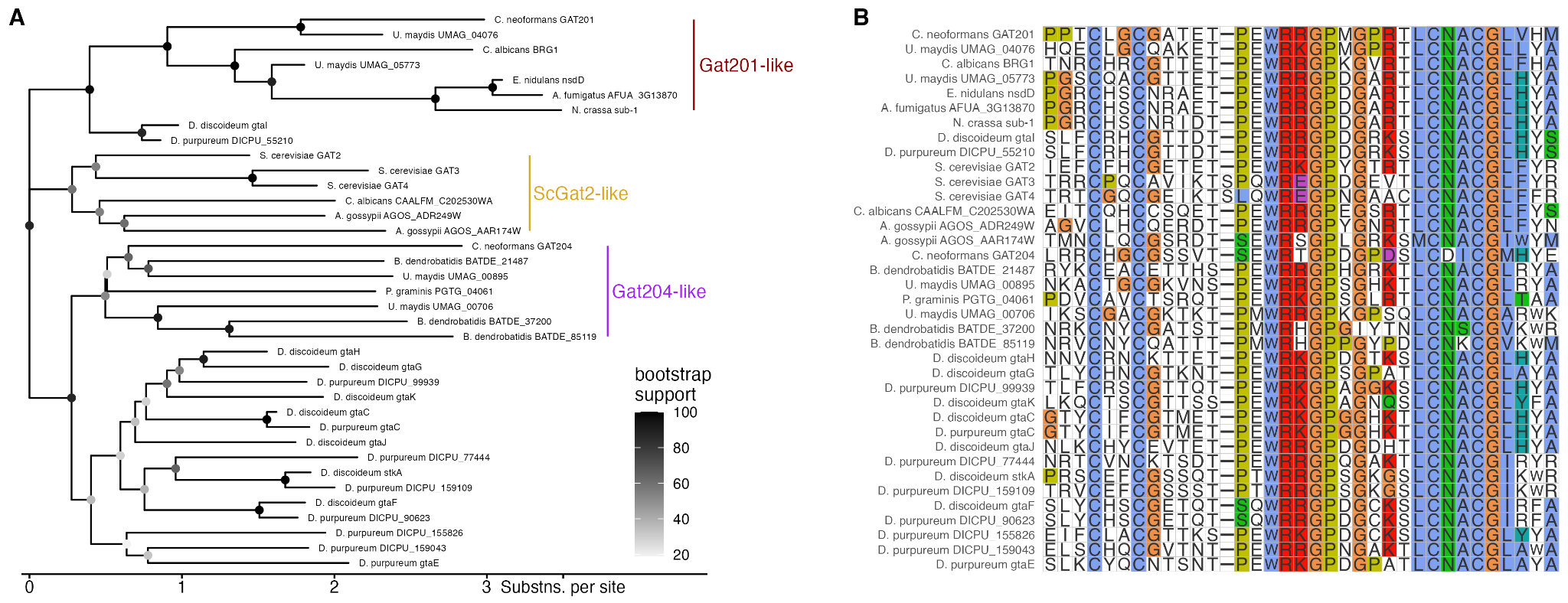
